# Systematic mass-spectrometry-guided metabolic fingerprinting elucidates diversity of specialized metabolites across the Brassicaceae

**DOI:** 10.64898/2026.04.17.719190

**Authors:** Felicia C. Wolters, Tina Woldu, M. Eric Schranz, Marnix H. Medema, Klaas Bouwmeester, Justin J. J. van der Hooft

## Abstract

- Plants produce diverse bouquets of specialized metabolites (SMs), yet only a fraction of the vast phytochemical space has been explored to date. Comparative analysis of SM profiles can reveal hotspots of biochemical novelty, while systematic profiling across taxonomic levels does presently not cover large plant families.
- To study core and accessory SM profiles in the Brassicaceae plant family, we fingerprinted 14 species by Liquid-Chromatography Mass-Spectrometry (LCMS/MS). We develop standardized experimental and computational workflows integrating *in silico* annotation tools to study consensus compound class and substructure distributions of SMs. Furthermore, we investigate the congruence of chemotaxonomy and species phylogeny across an extended panel of 17 species.
- Unique metabolite profiles were outstanding in *Camelina sativa, Capsella rubella*, and *B. vulgaris*, with the largest unique terpenoid profile annotated in *C. sativa*, accounting for 33.5% and 55.6% in positive and negative ionization mode, respectively. Substructure motifs were found to overlap with compound class predictions, highlighted for triterpenoids in Camelinodae. Furthermore, dual-tissue chemotaxonomic clustering resembled relationships of *Brassica* subgenomes across tissues.
- We anticipate that our systematic approach can serve as a blueprint for investigating biochemical diversity in other plant lineages and can boost the characterization of plant natural product pathways.

## Introduction

Plants rely on a vast array of chemical defenses to compensate for their sessile nature, representing one of the most structurally diverse and largely unexplored metabolite suites. Throughout human history, the rich chemical space of plant specialized metabolites (SMs) has been explored for its potential applications in pharmacy and food industry, with numerous examples of plant-derived drugs (Wang *et al*., 2019), food additives (Costa-Pérez *et al*., 2023), and applications for crop resistance breeding (Afifah *et al*., 2019; Macel *et al*., 2019; Tinte *et al*., 2021). Our understanding of chemodiversity in plants has grown in scale with the aid of novel computational tools, utilizing the increasing amount of high-resolution data from Gas and Liquid Chromatography tandem Mass Spectrometry (GC- and LC-MS/MS) platforms. Depending on the methods used for compound extractions (e.g. choice of solvents, gradients, instrument types), metabolic profiling captures structural diversity under positive and negative ionization modes at high resolution. In recent years, metabolic fingerprinting studies explored the metabolite diversity across multiple taxonomic levels, including plant family (Kang *et al*., 2019; Liu *et al*. 2021), genus (Elser *et al*., 2023) and species (Bakir *et al*., 2020; Fiehn *et al*., 2000; Houshyani *et al*., 2012; Padilla-González *et al*., 2023; Qiu *et al*., 2024) on an extensive scale. Intraspecific SM profiles were captured across genotypes (Tohge *et al*., 2018), individual plants (Mönchgesang *et al*., 2016) and tissues, including leaf and root tissue (Marzouk *et al*., 2023), seeds and fruits (Anaia *et al*., 2025; Barreda *et al*., 2024, 2025; Farag *et al*., 2024; Nguyen *et al*., 2025).

Plant adaptation to ecological niches is further differentiated by ontogeny and biotic and abiotic factors, with specialized biochemistry co-evolving with herbivores, pathogens, and microbiomes (Anaia *et al*., 2025; Goodger *et al*., 2013; McLaughlin *et al*., 2023). Yet, research focusing on metabolic fingerprinting does often not incorporate dimensions of tissues across species taxonomy within the same study. Consequently, the systematic study of chemical diversity across plant evolutionary history is still in its infancy, partially due to limitations in reliable compound annotation and integration of LC-MS/MS data across platforms, experiments, and batches (Alseekh and Fernie 2023; Houriet *et al*., 2025).

In recent years, the options for large-scale and pan-repository metabolomics data integration have been facilitated by the advancement of computational strategies. Nevertheless, metabolomics data integration across batches and experiments is still hampered by shifts in retention time (RT) and differences in instrument settings, interfering with the alignment based on RT windows and fragmentation patterns (Hajjar *et al*., 2023). Beyond sensitivity to technical settings in LC-MS/MS analysis, metabolic fingerprints are determined by biological variation during plant development, and by environmental conditions, underscoring the need for standardization of experimental setups. Hence, the systematic collection of LC-MS/MS data under standardized conditions is a prerequisite for robust comparative analysis, and identification of chemodiversity hotspots (Wolters *et al*., 2024). Contextualization of comparative SM profile analysis across plant taxonomy further relies on the resolution of species phylogeny. Plant families such as the Brassicaceae (crucifers) have been studied extensively to model evolutionary trajectories of genomic rearrangements after polyploidization (German *et al*., 2023; Hendriks *et al*., 2022; Walden & Schranz, 2023).

The Brassicaceae family is known for its economically important species cultivated as vegetable and oil crops, ornamentals, and the model species *Arabidopsis thaliana*. Recently, a comprehensive phylogeny of the Brassicaceae family has been released. Together with a high-resolution phylogeny, Hendriks *et al*. (2024) introduced a revised family classification, covering two subfamilies (Brassicoideae and Aethionemoideae) and five supertribes (Camelinodae, Brassicodae, Arabodae, Heliophilodae, Hesperodae). The evolution of glucosinolates in the Brassicaceae family as a characteristic SM class has been studied extensively (Agerbirk *et al*., 2021; Bird *et al*., 2025). Other compound classes such as triterpenoid saponins were exclusively found in one genus (*Barbarea*) in the Brassicaceae (Erthmann *et al*., 2018; Li *et al*., 2023). Yet, untargeted metabolic fingerprinting studies do presently not cover multiple related species across supertribes in the Brassicaceae family.

Here, we present a systematic comparative analysis of SM class distributions across the Brassicaceae, focusing on the supertribes Camelinodae and Brassicodae. To investigate shared and unique SM profiles, we collected and analyzed LC-MS/MS data from 14 Brassicaceae species and two tissues (leaf and root), incorporating feature extraction in mzmine, computational annotation of extracted metabolite features, and feature-based molecular networking (FBMN) in both positive and negative ionization mode. We specifically highlight the shared metabolic profile - hereafter referred to as “core-metabolome” - across supertribes, genera and species, and in both leaf and root tissue. In addition, we investigate the distribution of alkaloid and terpenoid SM classes across taxonomic levels and tissues, and examine the distribution of substructures corresponding to oleanane triterpenoid compound class annotations. Ultimately, we compared chemotaxonomic relationships based on hierarchical clustering with species phylogeny across an extended set of 17 species, to determine congruency across tissue-specific metabolite profiles.

## Materials and methods

### Plant material and growth conditions

Seeds of plant genotypes listed in Table 1 were vapor-sterilized using chlorine gas in a desiccator for 3h, cold stratified at 8°C for seven days, and sown on half-strength Murashige and Skoog (MS) agar. Plants were grown for 25 to 35 days in a climate chamber (16:8h photoperiod, 140-180 µmol m2 s-1, 22 ± 2°C) until seedlings developed four mature leaves (PO:0007115). Plant positions in the climate chamber were randomized.

**Table 1.** Used plant species and genotypes organized by taxonomic order.

| **Supertribe<sup>a</sup>** | **Tribe<sup>a</sup>** | **Genus** | **Species** | **Accession code** |
| --- | --- | --- | --- | --- |
| Brassicodae | Brassiceae | *Brassica (Mutarda)* | *nigra*<sup>1</sup> | SF48 |
| Brassicodae | Brassiceae | *Hirschfeldia* | *incana*<sup>1</sup> | Nijmegen |
| Brassicodae | Brassiceae | *Eruca* | *vesicaria*<sup>1</sup> | PI 6333218 |
| Brassicodae | Brassiceae | *Brassica* | *carinata*<sup>1</sup> | Gomenzer |
| Brassicodae | Brassiceae | *Brassica* | *juncea*<sup>2</sup> | CGN15183 |
| Brassicodae | Brassiceae | *Brassica* | *oleracea*<sup>1</sup> | CGN18947 |
| Brassicodae | Brassiceae | *Sinapis* | *alba*<sup>1</sup> | S2 GC0560-79 |
| Brassicodae | Isatidae | *Isatis* | *tinctoria*<sup>1</sup> | Weber |
| Camelinodae | Camelineae I | *Camelina* | *sativa*<sup>1</sup> | Daan-15 |
| Camelinodae | Camelineae I | *Capsella* | *rubella*<sup>1</sup> | Monte Gargano |
| Camelinodae | Arabidopsideae | *Arabidopsis* | *thaliana*<sup>1</sup> | Col-0 |
| Camelinodae | Lepideae | *Lepidium* | *sativum*<sup>1</sup> | SluisG |
| Camelinodae | Cardamineae | *Barbarea* | *vulgaris*<sup>1</sup> | Zodiac |
| Camelinodae | Cardamineae | *Rorippa* | *islandica*<sup>2</sup> | GC 2006-89 |
| Camelinodae | Cardamineae | *Cardamine* | *hirsuta*<sup>1</sup> | De Hoef |
| Arabodae | Arabideae | *Arabis* | *alpina*<sup>2</sup> | Pajares |
| Outgroup | Aethionemeae | *Aethionema* | *arabicum*<sup>1</sup> | ES1020 |

<sup>1</sup>Included in batch A (14 species); <sup>2</sup>Included in batch B (3 species); <sup>a</sup>Phylogeny according to Hendriks *et al.* (2023)<sup>5</sup>.

### Plant tissue sampling

Plants were harvested from agar pots, and leaf and root tissue were carefully separated, using a scalpel, excluding stem and cotyledons. Residual growth medium was removed from roots by washing with demineralized water. Harvesting was performed after 8-10 h of light exposure (between 1 and 3 PM), and collected tissues were immediately frozen in liquid nitrogen. Samples from three biological replicates per species consisting of 5-20 plants per replicate grown in independent tissue culture units were harvested and homogenized under liquid nitrogen using a mortar and pestle and stored at −80°C until further analysis.

### Metabolite extraction and LC-MS/MS analysis

Tissue samples were processed for metabolite extraction and analysis according to De Vos *et al*. (2007). Briefly, solvent (99.875% methanol acidified with 1.25% formic acid) was added to fresh tissue material in a 1:3 ratio (fresh weight tissue to solvent), followed by sonication for 15 min at 40 kHz in a water bath at 20°C. The liquid phase was obtained after centrifugation at 15,000 g. Samples were randomly injected (volume: 10 µl) at 20°C into a Vanquish UHPLC with Exploris120 Orbitrap system (Thermo Fisher Scientific, Waltham, MA, USA). The used Vanquish UHPLC was equipped with an Vanquish photodiode array detector (220–600 nm) connected to an Orbitrap Exploris 120 mass spectrometer fitted with an electrospray ionization (ESI) source. A reverse-phase column (Acquity UPLC BEH C18, 1.7 μm, 2.1 × 150 mm; Waters) was used at 40°C for chromatographic separation. Samples were analyzed with a flow rate of 0.4 ml/min with stepwise peak ratios of mobile phase eluents A (0.1% formic acid in ultrapure water), and B (0.1% formic acid in acetonitrile) with initial conditions set to 95:5 (A:B). After the linear gradient of 22 minutes (0.0 – 22.0 min, 5-75% B), washout phases (1min gradient 75% to 90% B, 3 min at 90% B) were followed by recalibration to initial conditions (95%A:5%B). Scans were obtained with an m/z range of 90-1350 Da in positive and negative mode (switched polarity) and a resolution of 60,000 FWMH. Data was obtained in DDA mode (centroid, Number of Dependent Scans = 4) with normalized HCD Collision Energies (%) = 20,40,100. Mass exclusion lists (Material S1) were generated from solvent blank samples injected prior to the tissue sample extracts, to avoid fragmentation of mass features present in the solvent under DDA mode. Quality Control (QC) samples were prepared from pooled samples, including material from all analyzed samples.

**Figure 1.**
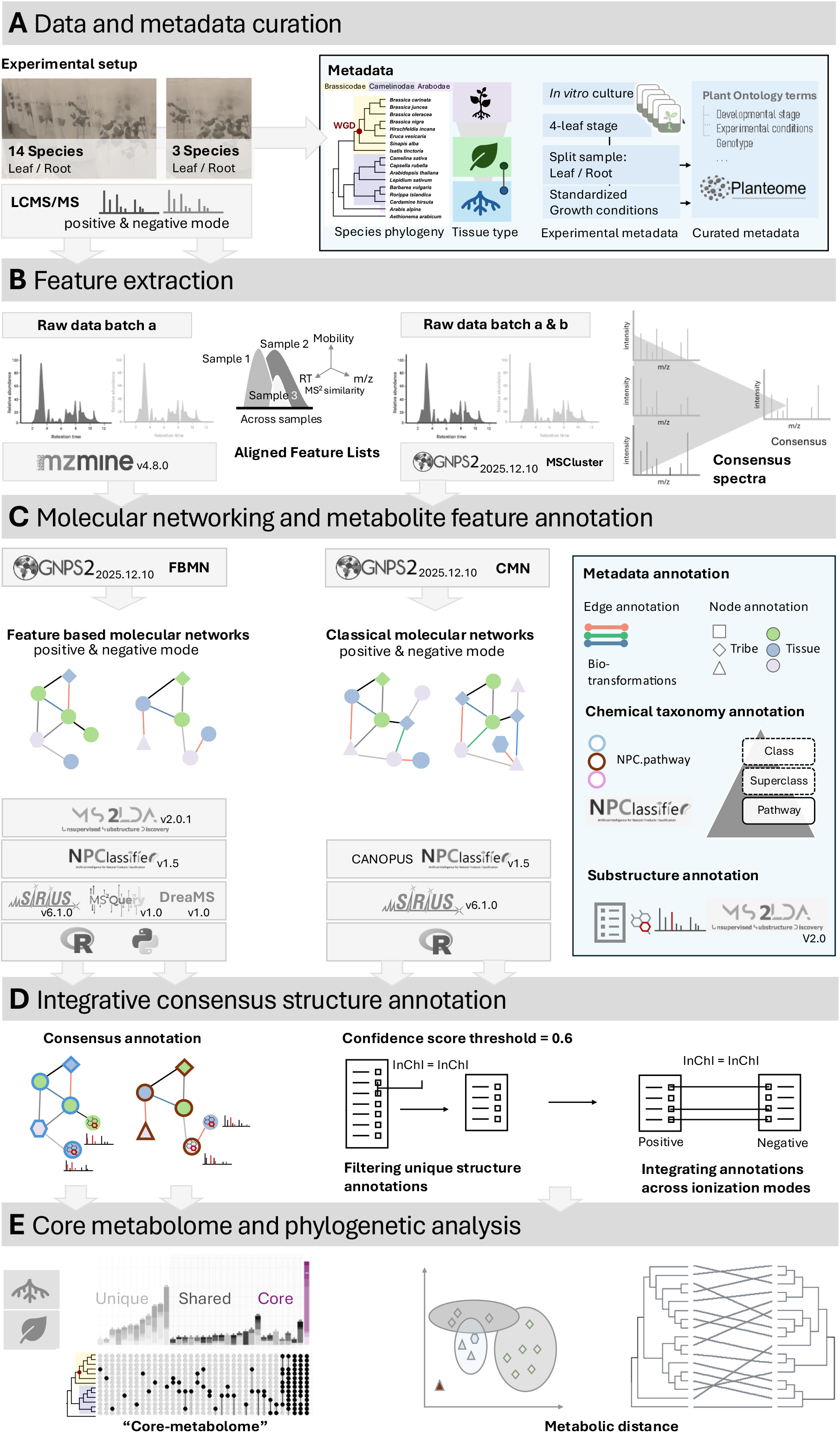
Workflow for untargeted metabolomics data collection, metadata curation, compound annotation, and batch integration for chemical taxonomy analysis. **A)** Data and metadata curation. Batches were analyzed independently on the LC-MS/MS platform using the same protocol. **B)** Feature extraction was conducted in mzmine for batch “a” including 14 species. Converted MZXML files of batch “a” and “b” including in total 17 species were submitted to the GNPS2 online platform, utilizing MSCluster for clustering spectra based on cosine similarity. **C)** Molecular networking and metabolite feature annotation. Batch “a” and combined batches “a” and “b” were submitted for Feature-Based Molecular Networking (FBMN) and Classical Molecular Networking (CMN) to the GNPS2 platform, respectively. The FBMN was annotated using SIRIUS including CANOPUS, MS2Query, and DreaMS, and chemical ontology annotations were obtained from NP.Classifier. Batch “b” was annotated using SIRIUS with a cutoff confidence exact score of 0.6. **D)** Integrative and consensus structure annotations were obtained by matching structural and chemical ontology predictions separately for annotation tools in batch “a”. In batch “b”, similar structure predictions were merged across multiple features, and across ionization modes. **E)** Core metabolome and phylogenetic analysis were conducted for the datasets including 14 and 17 species, respectively.

### LC-MS/MS data processing

Raw Waters MS data was converted to .*mzXML* file format using MSConvert v3.0. Extraction of metabolite features was conducted in mzmine v4.8.0 (Schmid *et al*., 2023) for a Retention Time (RT) window between 0.8 and 22.8 min. Presets are provided as .*mzbatch* file in XML format (Material S2). Aligned feature lists generated in mzmine were exported in MGF spectra format, and feature quantification tables in .csv format. Feature lists generated for spectral data of 14 species, and mzXML files of the combined dataset including spectral data of 17 species were submitted to the Global Natural Products Social (GNPS) Molecular Networking online platform using the Feature-Based Molecular Networking (FBMN) and Classical Molecular Networking (CMN) workflows, respectively (GNPS2 Release 2025.12.10, Wang *et al*., 2016).

### Structure and compound class annotation

Extracted features of 14 species were annotated in MS2Query v1.0 (De Jonge *et al*., 2023), DreaMS v1.0 (Bushuiev *et al*., 2025), and in SIRIUS v6.1.0 (Dührkop *et al*., 2021a), including assessment of prediction accuracy in ZODIAC (Ludwig *et al*., 2019), and structure annotation by CSI:FingerID (Dührkop *et al*., 2015). Spectral library searches using DreaMS v1.0 utilized the MassSpecGym spectral library (Bushuiev *et al*., 2024). Compound class annotations based on NP.Classifier (Kim *et al*., 2020) were obtained in CANOPUS (Dührkop *et al*., 2021b) for fingerprints computed in SIRIUS v6.1.0, and for analog predictions in MS2Query v1.0. NP.Classifier annotations were computed via the API (v1.5) for all structure matches and analog predictions obtained with DreaMS v1.0 regardless of similarity scores. Consensus structure and chemical ontology predictions were computed separately for exact matches of InChI key and compound class strings obtained in SIRIUS v6.1.0 and CANOPUS, and converted from SMILES obtained in DreaMS v1.0 and MS2Query v1.0, using an in-house script available on GitHub (https://github.com/capsicumbaccatum/Manuscript_Metabolic_profiling).

Total number, compound class categories and overlap of annotations per annotation tool are available on Zenodo (DOI: 10.5281/zenodo.19391098) and visualized in Supplementary Fig. S1. Combined output of metabolite feature annotations and consensus predictions was mapped on the FBMN. The consensus spectra of the CMN obtained for the combined dataset including 17 species was annotated with SIRIUS v6.1.0, using a cutoff of 0.6 for the Exact Confidence Score in downstream analysis.

### Mass2Motif analysis

Mass2motifs were obtained with MS2LDA v2.0.1 (van der Hooft *et al*., 2016; Wandy *et al*., 2017; Torres-Ortega *et al*., 2025), with settings provided in Material S3. In short, the number of discovered motifs was set to 399, and the number of iterations to 1000, using default settings for all other parameters. Mass2motifs were then mapped onto the FBMN with a doc-topic probability score threshold of 0.5 and an overlap score threshold of 0.15.

### Hierarchical clustering, PCA and chemical taxonomy analysis

Hierarchical clustering was computed for a distance matrix generated by the *dist()* function, specifying Euclidean distance. The *hclust* object was computed based on complete clustering (R package *stats* v4.5.1, R Core team, 2025) and subsequently converted to a dendrogram object for plotting against species phylogeny (according to Hendriks *et al*., 2024) using the *tanglegram* function of the R package *dendextend* v1.19.1. Tanglegrams were constructed for the full set of features in the CMN consensus MGF, for a subset of annotated features using SIRIUS v6.0.1 structure predictions with a Confidence Exact Score threshold of 0.6, and for features specific to leaf and root tissue, respectively. Based on the alignment with species phylogeny, tanglegrams are presented for comparative analysis between tissue types.

### Data visualization and formatting

Analyses and visualizations conducted in R v4.5.1 (R Core team, 2025) employed the following packages: DPLYR (Wickham et al. 2014) and TIDYVERSE (Wickham *et al*., 2019) for data formatting and filtering; GGPLOT2 (Schloerke *et al*., 2010), GGTREE and GGTREEXTRA (Xu *et al*., 2021; 2022), COMPLEXUPSET (Krassowski *et al*., 2022) and VENN (Ruskey & Weston 2005) for visualization and annotation of phylogenetic trees and upset plots. Analyses and visualizations conducted in Python v3.12.0 employed the following packages: RDKit v2025_09_3 (Landrum *et al*., 2025) for SMILES string conversion and Matchms (Huber *et al*., 2020) and MS2LDA 2.0 (Ortega *et al*., 2025) for visualization of mass spectra, motif extraction and formatting.

## Results and discussion

### The “core-metabolome” reflects taxonomic relationships in the Brassicaceae

To study the composition of conserved metabolic fingerprints in the Brassicaceae family, mass features extracted from LC-MS/MS data were aligned across 14 species in the Camelinodae and Brassicodae supertribes, and subsequently annotated with chemical class terms according to the NP.Classifier chemical ontology (Kim *et al*., 2020). Mass features represent detectable ions of compounds in a complex mixture. In total, 7,683 and 6,433 mass features from raw LC-MS/MS spectra obtained from leaf and root tissue were extracted and aligned with mzmine in positive and negative ionization mode, respectively. Filtering for consistent NP.Classifier pathway annotations across annotation tools (SIRIUS, DreaMS against the MassSpecGym library, MS2Query) resulted in 5,118 features in positive ionization mode (Fig. 2a), and 2,542 features in negative ionization mode (Fig. 2b). Alkaloids, amino acids and peptides, carbohydrates, and fatty acids pathway annotations were predominantly found in positive ionization mode across all species (Fig. 2a). In contrast, terpenoids, alkaloids, shikimates and phenylpropanoids were disproportionally enriched in profiles under negative ionization mode (Fig. 3b).

**Figure 2.**
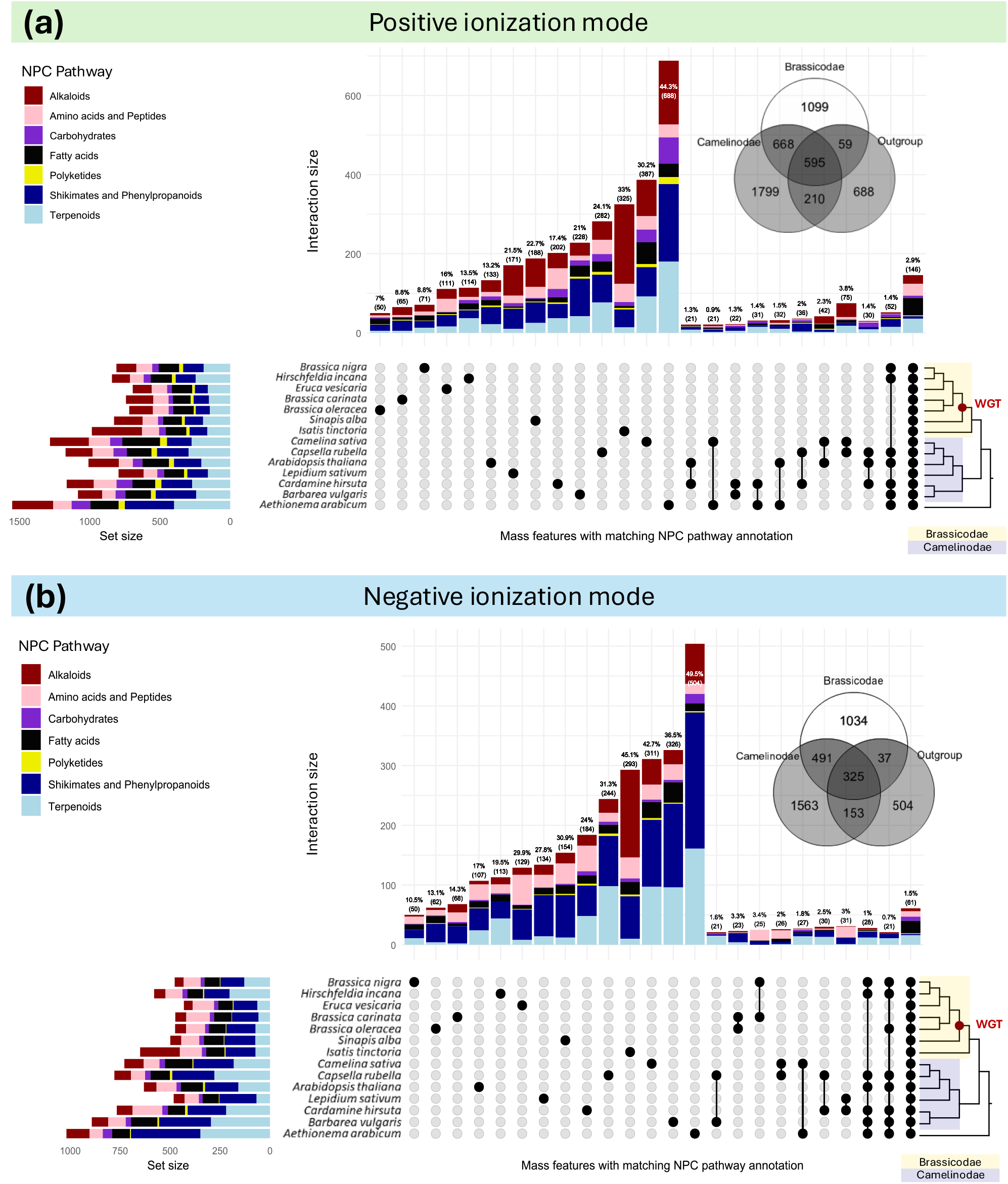
Distribution of mass features across 14 Brassicaceae species under positive and negative ionization mode. Untargeted metabolomics (LC-MS/MS) data was filtered for consistent annotations using computational annotation tools (SIRIUS, MS2Query, DreaMS) on the NP Classifier pathway chemical ontology level. Red circles on phylogenetic trees indicate the Whole Genome Triplication (WGT) event in the Brassicodae supertribe. Venn diagrams show overlaps of aligned features across supertribes. Only sets with at least 20 metabolite features are shown. **A:** Metabolite features annotated in positive ionization mode. **B:** Metabolite features annotated in negative ionization mode.

**Figure 3.**
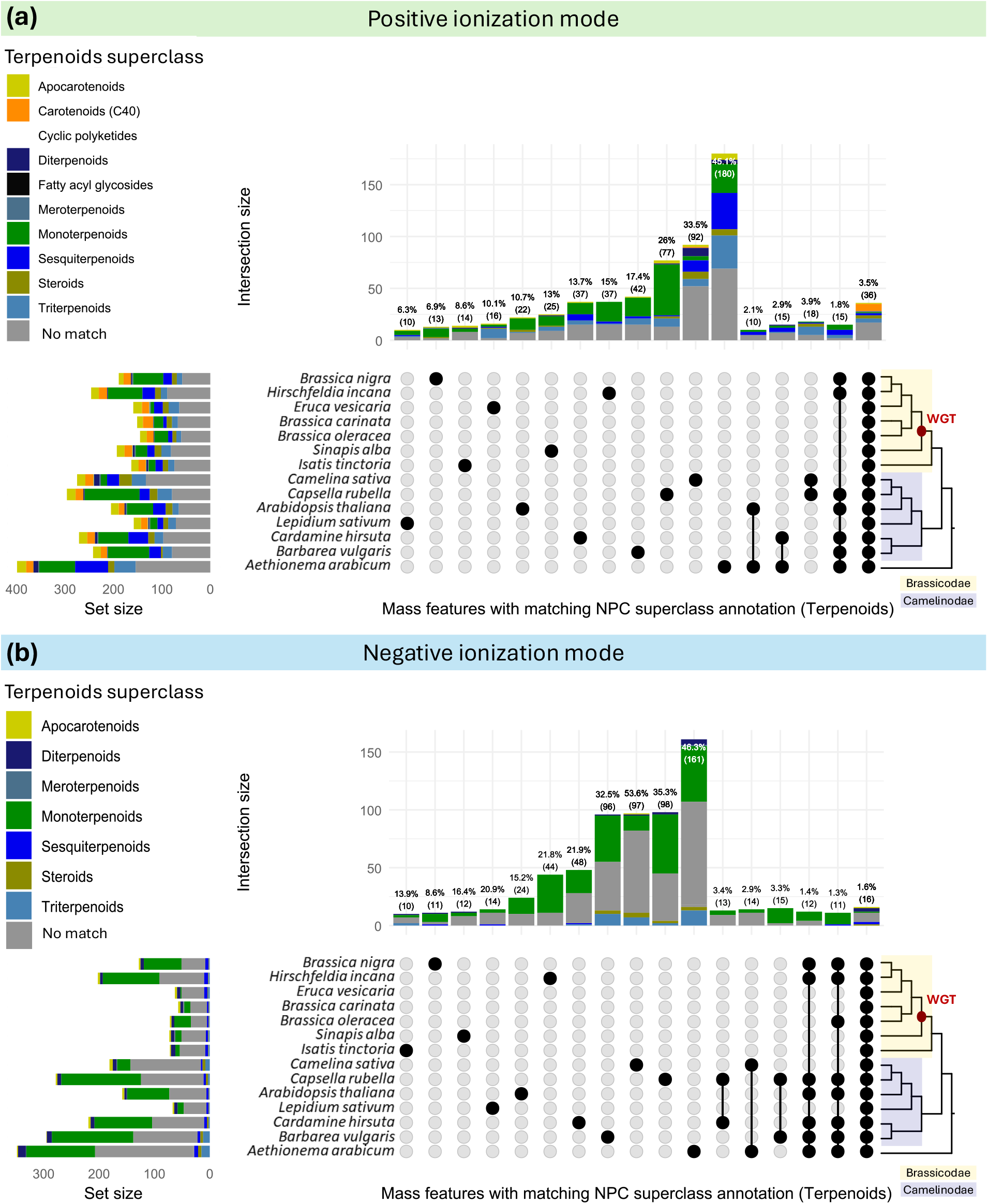
Distribution of mass features across 14 Brassicaceae species under positive and negative ionization mode. Untargeted metabolomics (LC-MS/MS) data was filtered for consistent annotations using computational annotation tools (SIRIUS, MS2Query, DreaMS) on the NP.Classifier superclass chemical ontology level, filtered for “terpenoid” pathway annotations. Terpenoid pathway annotations with inconsistent superclass annotation across at least two annotation tools are indicated in grey (“no match”). The red dots placed on the phylogenetic tree indicate the Whole Genome Triplication (WGT) event in the Brassicodae clade. Venn diagrams show overlaps of aligned features across supertribes. **A:** Metabolite features annotated in positive ionization mode. **B:** Metabolite features annotated in negative ionization mode.

The “core” set of annotated metabolites in the Brassicaceae shared by at least one species within Camelinodae, Brassicodae, and the outgroup is represented by 595 (11.6%) metabolite features in positive ionization mode, and 325 (12.8%) features in negative ionization mode (Fig. 2a). Brassicodae and Camelinodae share 668 (13%) metabolite features in positive ionization mode (Fig. 2a), and 491 (19.2%) features in negative ionization mode (Fig. 2b), representing the “core metabolome” shared by both supertribes. The unique metabolite profile of the outgroup species *Aethionema arabicum* is comparable in size with the unique profiles of both supertribes, with 13% in positive and 19.8% in negative ionization mode, suggesting balanced representation of both supertribes and the outgroup. A smaller fraction of annotated metabolite features is conserved on the species level, with 2.9% (146) and 1.5% (61) of all consistently annotated features shared across all species in positive and negative ionization mode, respectively. The shared sets of metabolite features across species are characterized by annotations for primary metabolite classes, such as amino acids and peptides, carbohydrates, and fatty acids. Within the Camelinodae supertribe, species share in total more than 200 features, predominantly annotated as terpenoids, alkaloids or shikimates and phenylpropanoids in positive ionization mode. The number of shared metabolite annotations across sets of species within the Camelinodae reflects taxonomic relationships, with *Camelina sativa* and *Capsella rubella* sharing 75 (3.8%) annotated features, followed by *C. sativa* and *Arabidopsis thaliana* (42, 2.3%) in positive ionization mode (Fig. 2a). In contrast, species of the Brassicodae supertribe share smaller sets with less than 20 features (2%) under positive ionization mode (Fig. 2a). In negative ionization mode, *Brassica carinata* shares 23 (3.3%) features primarily annotated as amino acids and peptides with *Brassica nigra*, and 25 (3.4%) features with *Brassica oleracea*, mostly annotated as shikimates and phenylpropanoids (Fig. 2b). All other species of the Brassicodae supertribe have less than 20 metabolite features with consistent NPC.pathway annotations in common (Material S6).

Our findings suggest that conserved profiles across supertribes reflect phylogenetic distance, highlighted by the overlaps of Camelinodae, Brassicodae and outgroup in both ionization modes. However, unique metabolite features represent a broad range of 7% to 45% of the total captured fingerprint, especially pronounced in the Camelinodae and under negative ionization mode. The higher numbers of species-specific detected and consistently annotated metabolite features in Camelinodae compared with Brassicodae can only partly be attributed to the larger overall feature set observed in Camelinodae species. Notably, species representing the Camelinodae supertribe also cover larger phylogenetic distances compared to Brassicodae species. The represented tribes within the Camelinodae supertribe cover Isatidae, Camelinae I, Arabidopsidae, Lepidae, and Cardaminae, while the Brassicodae supertribe is mainly limited to the Brassiceae tribe. Recent meta-analyses indicate that, although the discovery of new plant compounds shows signs of saturation as more species are studied, the known chemical space still represents only a small fraction of plant diversity (Domingo-Fernández *et al*., 2023; Hart *et al*., 2025). To better estimate undiscovered chemical diversity and reveal shared biochemical space, we propose systematic metabolomic fingerprinting across more taxonomic levels within the Camelinodae supertribe.

### Metabolite profiles of *Camelina sativa, Isatis tinctoria, Capsella rubella*, and *Barbarea vulgaris* contain large numbers of unique features

To investigate the size and composition of unique metabolite profiles across species, we compared the proportions of annotated compound classes in the “accessory” (unique) metabolome. In total, between 7% and 49.5% of detected metabolite features were found to be unique per species. Across all species, unique metabolite profile annotations were enriched with alkaloids (27.96%), shikimates and phenylpropanoids (25.54%), and terpenoids (19.07%) in positive ionization mode. In negative ionization mode, enrichment with shikimates and phenylpropanoids (37.92%), and with terpenoids (23.48%) was found higher than with alkaloids (15.23%) in annotated unique metabolite profiles. Across both ionization modes, the outgroup species *A. arabicum* exhibited the highest number of unique features with matching NP.Classifier annotations, with 688 (44.3%) metabolite features in positive ionization mode, and 504 (49.5%) metabolite features in negative ionization mode (Fig. 2a & 2b). In positive ionization mode, *Camelina sativa* showed the highest total number of unique mass features (387, 30.2%) within the Camelinodae, followed by *Isatis tinctoria* (325, 33%) in the Brassicodae. Notably, *I. tinctoria* (Isatidae tribe) also possessed the largest number of unique alkaloid pathway annotations across all intersections and in both ionization modes, resembling taxonomic distance to species in the Brassiceae sister tribe (Fig. 2a & 2b). In negative ionization mode, *Barbarea vulgaris* displayed the largest unique profile of annotated mass features (326, 36.5%) in the Camelinodae. However, corrected for the total set size (total number of detected metabolite features) per species, *I. tinctoria* showed the largest proportion of unique features in positive (33%) and negative (45.1%) ionization mode (Fig. 2b). In summary, the accessory metabolite profiles of three species in the Camelinodae supertribe (*B. vulgaris, C. sativa, C. rubella*) and in *I. tinctoria* (Brassicodae) are outstanding, with terpenoid profiles disproportionally enriched in Camelinodae species.

The outstanding unique metabolite profiles of *I. tinctoria* and *B. vulgaris* were found in alignment with previous research focusing particularly on alkaloid and terpenoid diversity in *I. tinctoria (Xu et al*., *2020)* and *B. vulgaris* (Erthmann *et al*., 2018; Shen *et al*., 2025; Shinoda *et al*., 2002), respectively. Furthermore, our results suggest that the metabolite profile of *C. sativa* is particularly distinctive, also representing the largest terpenoid profile next to the outgroup *A. arabicum* in positive ionization mode (Fig. 3a). The evolutionary history of *C. sativa* may partially explain its outstanding unique metabolite profile. *C. sativa* has undergone several rearrangements of its hexaploid genome (*n*=20), which arose from hybridization between a *C. neglecta*-like genome (*n*=6) and *C. hispida* (*n*=7) (Mach *et al*., 2019). The characterization of the metabolite profile of *C. sativa* has been mostly focused on mono- and sesquiterpenoids (Augustin *et al*., 2019), amino acids, flavonols, cinnamic acids and glucosinolates in previous research (Alberghini *et al*., 2022; Guendouz *et al*., 2022; Zhang & Parkin, 2025). Our findings suggest that the terpenoid profile of *C. sativa* constitutes a diverse mixture of terpenoid classes, including triterpenoids, and is outstanding and unique in the panel of species included in our analysis. While the high content of polyamines in *C. sativa* has been linked earlier to advanced resilience against abiotic stress (Zhang & Parkin, 2025), terpenoid diversity has not yet been studied in this species in greater detail to our knowledge, especially with regards to high-mass compounds (> m/z 1000). The number of unique features in the terpenoid NPC.pathway consistently annotated in *C. sativa* (92, 33.5%) under positive ionization mode is about two-fold higher than in *B. vulgaris* (42, 17.4%), while the latter is known to produce a diverse range of specific terpenoids (Khakimov *et al*., 2016). Studying metabolite profiles of *C. sativa* progenitors (*C. neglecta* and *C. hispida*) may elucidate the impact of genome rearrangements following polyploidization in *C. sativa*, which has potentially resulted in the diversification of terpenoid biosynthesis.

### The unique profiles of species in the Camelinodae supertribe are characterized by triterpenoids

Terpenoid annotations were disproportionally enriched in unique profiles of species, specifically within the Camelinodae supertribe. The largest unique profiles of annotated terpenoids were comparable in size between *C. sativa* (92, 33.5%) and *C. rubella* (77, 26%), and comprised mostly monoterpenoid annotations in *C. rubella*, which were underrepresented in *C. sativa* (Fig. 3). To further explore this pattern, we investigated compound class annotations on the superclass level to determine the distribution of mono-, di-, sesqui- and triterpenoids across species and supertribes. We found that annotated terpenoid profiles were characterized by mono-, di- and triterpenoid superclasses across supertribes, with monoterpenoids dominating in negative ionization mode.

Fewer consensus NP.Classifier superclass and class annotations were obtained for features detected in negative mode (Fig. 3b), potentially attributed to missing annotations in DreaMS v1.0 being presently restricted to positive ionization mode analysis. Triterpenoid superclass annotations had particularly inconsistent annotations on the class ontology level (Fig. 4, Table S6 available on Zenodo), potentially related to molecule size and complexity in comparison with mono- and diterpenoids.

**Figure 4.**
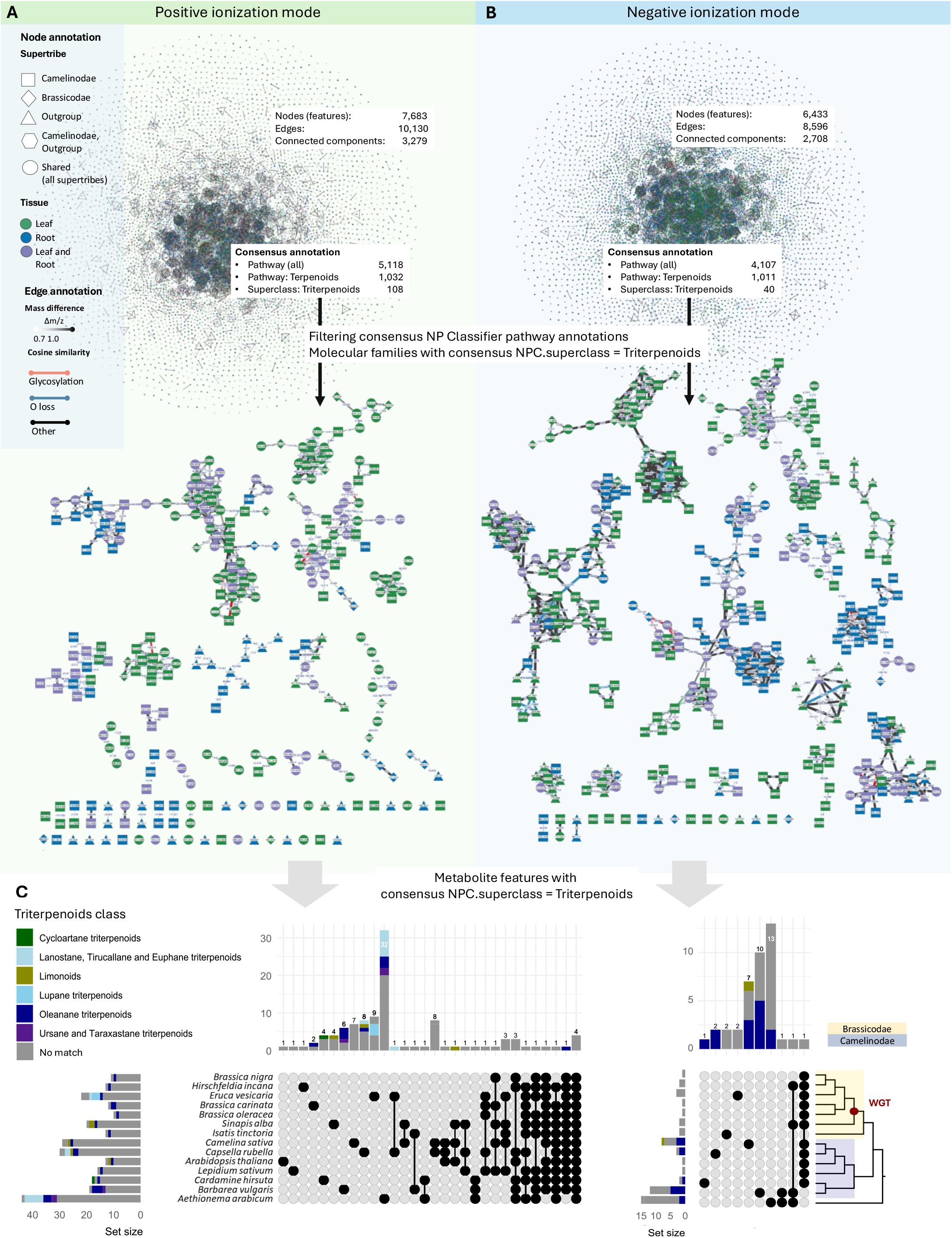
Distribution of triterpenoid consensus superclass annotations on the NP.Classifier superclass level. Feature-Based Molecular Networks (FBMNs) in (**A**) positive and (**B**) negative ionization mode. Node shapes represent presence/absence in supertribes, and node colors represent presence/absence in tissue types. Edges were annotated with biotransformations corresponding to glycosylation (162.05 Da, 146.06 Da, rhamnose; glucose or galactose; 176.03 Da, glucuronide; 133.05 Da, arabinoside; 324.1 Da, double glycosylation; 309.12 Da, rhamnose and glucose) and oxidation (15.995 Da). Consensus annotations of NP.Classifier superclass “triterpenoids” were filtered in both ionization modes. **C:** Upset plots showing distribution of NP.Classifier class annotations across species for metabolite features consistently annotated as “triterpenoids” on the superclass ontology level. Triterpenoid consensus annotations without a matching class annotation across at least two annotation tools are indicated in grey (“no match”). The red dot on the phylogenetic tree indicates the Whole Genome Triplication (WGT) event in the Brassicodae clade.

Consistently annotated mass features within the triterpenoid superclass belong to the classes of lupane, oleanane, lanostane, tirucallane, and euphane triterpenoids in positive ionization mode (Fig. 4a), whereas only putative oleanane-type triterpenoids and a single limonoid were annotated in negative ionization mode (Fig. 4b). Characteristic fragmentation patterns in MS2 under negative ionization mode may facilitate the annotation of oleanane-type triterpenoids, biasing compound class representation. The largest proportions of unique triterpenoid annotations were found in *A. arabicum, B. vulgaris*, and *C. sativa*. The unique triterpenoid profile of *B. vulgaris* consisted predominantly of putative oleanane triterpenoids in both ionization modes. In *C. sativa*, the unique triterpenoid profile comprised four consistent class annotations (putative oleanane triterpenoids and limonoids) in negative ionization mode (Fig. 4b), whilst the positive ionization mode did not result in any consistent class annotations (Fig. 4a). Thus, compound class predictions overlap with library matches obtained for previously described oleanane triterpenoid saponins in *B. vulgaris* (Khakimov *et al*., 2016; Shinoda *et al*., 2002). Furthermore, consistent annotations indicate the presence of putative oleanane triterpenoid saponins in other species, primarily in the Camelinodae supertribe.

### Substructures associated with oleanane triterpenoids and conjugated glycosides are not exclusive to *Barbarea vulgaris*

Presence of consensus class annotations in Camelinodae species indicated distribution of putative triterpenoid saponin-like metabolite features in other species apart from *B. vulgaris*. To investigate this further, molecular families including consistently annotated metabolite features were subjected to detailed analysis. To investigate the overlap of oleanane triterpenoid annotations with shared substructures, patterns of mass fragmentation and neutral losses (Mass2Motifs) were extracted using MS2LDA in both positive and negative ionization mode. Extracted Mass2Motifs were inspected with regards to substructure SMILES annotations and characteristic MS fragmentation patterns corresponding to oleanolic acid (Wikidata: Q418628) and conjugated triterpenoid saponin glycosides reported in the literature (Khakimov *et al*., 2016; Shinoda *et al*., 2002).

Three Mass2Motifs detected in positive ionization mode spectra were annotated as substructures of putative oleanane- and ursane-type triterpenoids. Motif_66 and motif_219 (Fig. S2) were found to be associated with an oleanolic acid substructure, whereas motif_110 matches conjugated oleanane triterpenoid glycosides (Fig. 5b). These motifs were detected across leaf and root tissue of five Camelinodae species, in *Brassica carinata* (Brassicodae), and in the outgroup *A. arabicum*. Pseudospectra of motif_66 and motif_219 resembled characteristic fragment peaks of oleanane-type triterpenoids (m/z 439.36, 119.09, 105.07) which were also found in library spectra of oleanolic acid (GNPS-Library accession: CCMSLIB00003140213) and referenced in the literature (Fig. 5a). Base peaks in the pseudospectrum of motif_66 at m/z 439, 437, 425 corresponded to triterpenoid saponin aglycones previously reported by Trugawa *et al*. (2019). Also, the distribution of motif_66 in metabolite features was found aligned with the triterpenoid superclass annotations from NP.Classifier, and with GNPS library matches for oleanolic acid and hederoside F (Wikidata: Q105301871) in *B. vulgaris*, which also share a consensus class annotation as putative oleanane-type triterpenoids. Another characteristic base peak (m/z 471) of oleanane-type triterpenoid saponins in *B. vulgaris* was detected in the MS1 spectra of *A. arabicum*. This peak occurred in two mass features with an analog match to betulonic acid methyl ester predicted by SIRIUS and MS2Query, and associated with motif_66 below the probability score threshold (probability score of 0.45; Table S6).

**Figure 5.**
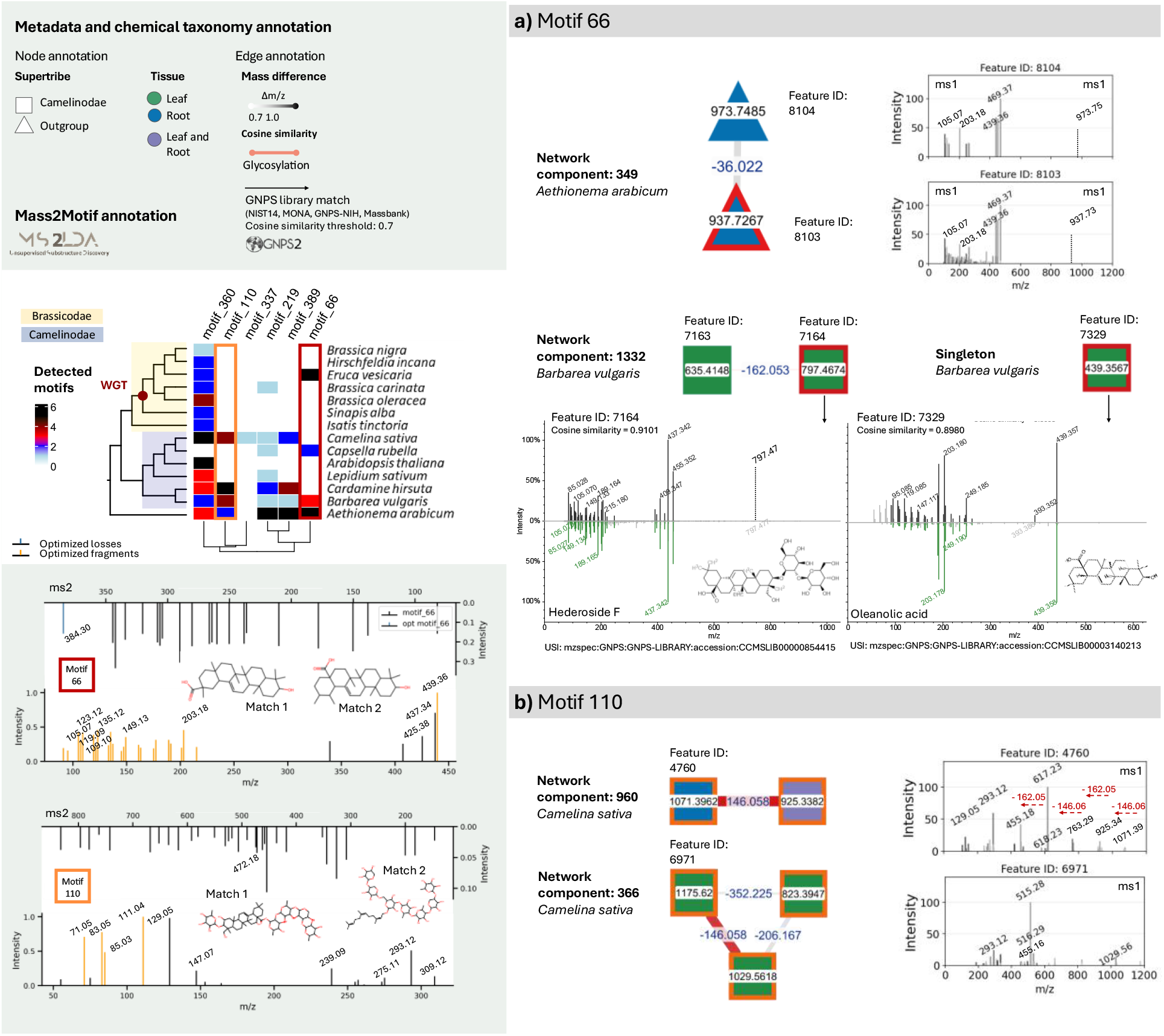
Mappings of Mass2Motifs extracted with MS2LDA in positive ionization mode. Mass2Motifs associated with fragmentation patterns and annotated substructures of oleanane-type triterpenoids were mapped on molecular families in the Feature-Based Molecular Network (FBMN). Node shape and color represent presence/absence in supertribes and tissues, respectively. Edges were annotated with biotransformations corresponding to glycosylation (162.05 Da, 146.06 Da) and hydroxylation (18.01 Da). **A:** Distribution of motif_66 and motif_219 across features with matching triterpenoid superclass prediction according to NP.Classifier chemical ontology, and library matches for spectra found in *Barbarea vulgaris*. **B:** Distribution of motif_110, and corresponding spectra of molecular features in *Camelina sativa, Cardamine hirsuta*, and *Aethionema arabicum*.

Motif_66 was extracted from spectra of 16 metabolite features (probability score >0.5, overlap score >0.15) in positive ionization mode. Of these, 14 features were consistently annotated as triterpenoids by NP.Classifier. Furthermore, motif_66 was found to overlap with GNPS library matches for previously described triterpenoid saponins in *B. vulgaris*. For two features annotated with motif_66 in *B. vulgaris*, spectral matches to oleanolic acid and hederoside F were obtained in the GNPS library. Notably, the aglycone peaks (m/z 437, 455) in the library spectrum of hederoside F overlap with motif_66 (m/z 437) and motif_219 (m/z 455), and are also shared by features annotated with motif_110 in *C. sativa* (Fig. 5a, Material S3). In addition, MS1 spectra of features sharing motif_110 exhibit characteristic putative aglycone peaks at m/z 445 and 455 that were not part of the MS2 motif. Another two features in *C. sativa* share hexoside (162.05 Da) and rhamnoside (146.06 Da) sugar losses in MS1 spectra (Fig. 5b). Optimized fragment peaks in motif_110 (m/z 71.05, 83.05, 85.03, 111.04) do not overlap with motif_66 and motif_219, but were annotated as fragments indicative of a glycosylation according to MotifDB (reference MotifSet: Massbank derived Mass2Motifs). Notably, molecular families annotated with motif_110 contained metabolite features with highly similar retention time, indicating in-source fragmentation (ISF). The impact of ISF on metabolite feature annotation in untargeted LC-MS/MS has been discussed recently (Abiedad *et al*., 2025). Therefore, putatively identified glycosylation patterns in network component 960 can potentially be attributed to ISF originating from the same molecule (Fig. 6b). Nonetheless, metabolite features prioritized by overlapping compound class and substructure annotations display characteristic fragmentation patterns of triterpenoid saponins that have not been reported in *C. sativa* and *A. arabicum* before.

**Figure 6.**
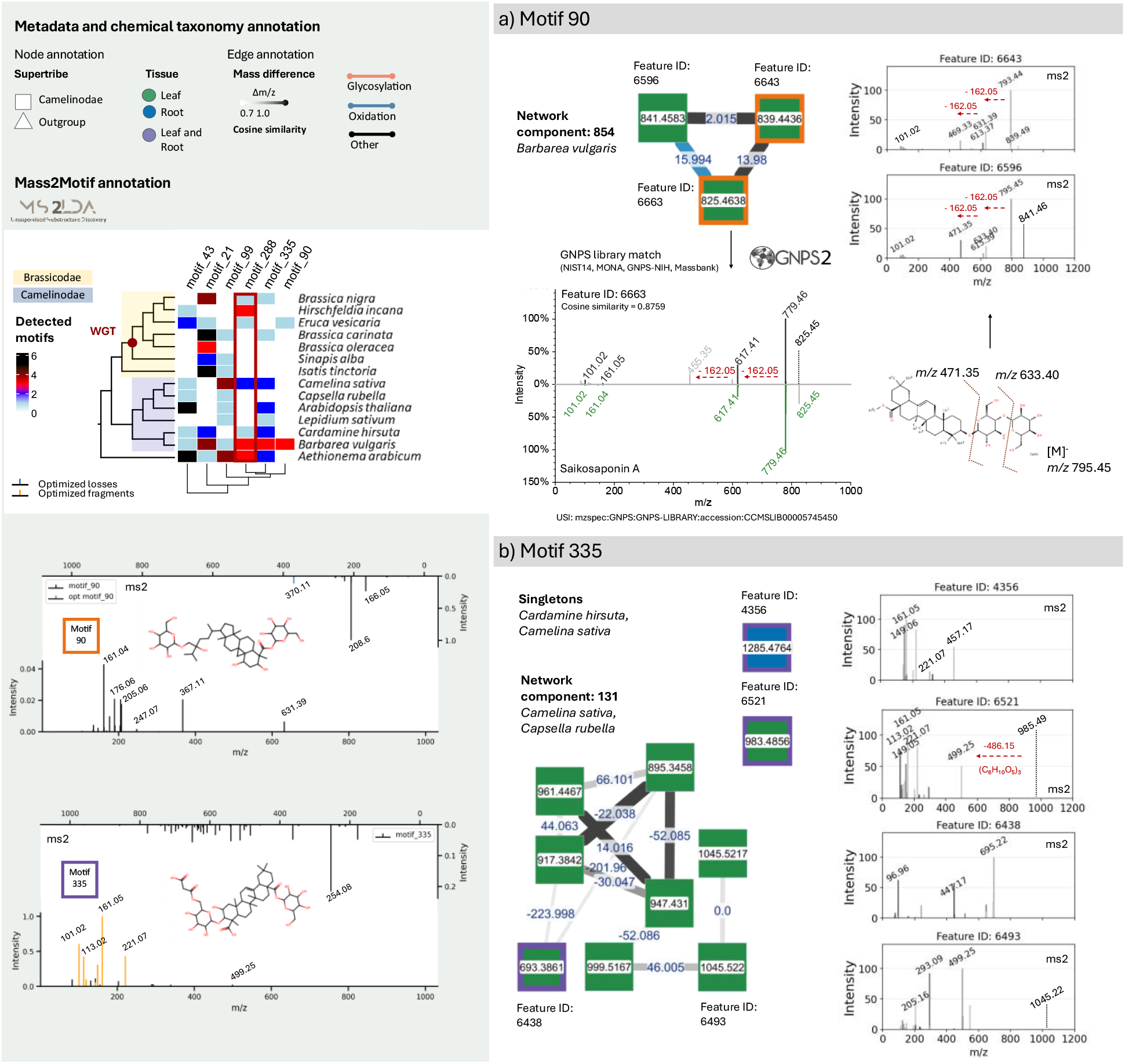
Mappings of Mass2Motifs extracted with MS2LDA in negative ionization mode. Mass2Motifs associated with fragmentation patterns and annotated substructures of oleanane-type triterpenoids were mapped on molecular families in the Feature-Based Molecular Network (FBMN). Node shape and color represent presence/absence in supertribes and tissues, respectively. Edges were annotated with biotransformations corresponding to glycosylation (162.05 Da, glucose) and oxidation (15.995 Da). **A:** Distribution of motif_90 across a selection of features, and GNPS library match in *Barbarea vulgaris*. **B:** Distribution of motif_288 on a selection of features in Camelinodae species.

In negative ionization mode, six pseudospectra of Mass2Motifs (i.e., motif_21, motif_43, motif_90, motif_99, motif_28, motif_335) were identified based on overlapping characteristic fragmentation patterns under negative ionization mode, and substructure SMILES annotations corresponding to oleanane-type triterpenoid saponin aglycones and conjugated glycosides (Fig. 6, Material S3). Motif_90, motif_335 and motif_288 were specifically distributed across features with matching superclass and class annotations, while motif_43, motif_21 and motif_99 were not associated with matching terpenoid consensus annotations on any chemical ontology level.

Motif_90 was extracted from a spectrum with library match to saikosaponin A (Wikidata: Q7499688) in *B. vulgaris*, a triterpenoid saponin previously characterized in a traditional Chinese medicine derived from extracts of *Bupleurum* spp. (Apiaceae; Yuan *et al*., 2017). However, the aglycone peak (m/z 455) and sugar losses were previously described by Khakimov *et al*. (2016) and Nielsen *et al*. (2010) in *B. vulgaris* G-type. Motif_90 is characterized by five mass peaks (m/z 161.04, 176.06, 205.06, 367.11, 631.39) corresponding to fragmentation patterns of triterpenoid saponin aglycones (Tsugawa *et al*., 2019), and two optimized losses (370.11 Da, 206.05 Da).

Distribution of motif_90 was found to be limited to leaf tissue profiles of *B. vulgaris* and *Eruca vesicaria*. In *B. vulgaris*, three aglycones (m/z 469, 471, 455) in network component 854, and corresponding sugar losses were tentatively identified as described in the literature (Khakimov *et al*., 2016; Shinoda *et al*., 2002; Fig. 6A). Motif_335 is shared by eight metabolite features with probability scores ≥0.5 and overlap scores ≥0.15, six of which were annotated as terpenoids by NP.Classifier (Table S6). While motif_355 is widely distributed across species in Camelinodae and Brassicodae, motif_90 is restricted to *B. vulgaris* and *E. vesicaria*. However, mass peaks (m/z 101.02 and 161.05) in matching library spectra (feature ID: 6663) were absent in motif_90, but present in motif_335 (Fig. 6b). In addition, most optimized fragment peaks of motif_335 (m/z 101.02, 113.02, 161.05, 221.07) overlap with motif_288 (Material S3). Motif_335 and motif_288 were found in leaf and root tissue across species of Camelinodae and Brassicodae supertribes, as well as in the outgroup. Five metabolite features (probability score ≥0.5, overlap score ≥0.2) share motif_288, four of which have a consensus annotation in the terpenoid pathway, while two share a triterpenoid superclass consensus annotation (Table S6). Metabolite features that were prioritized based on compound class and substructure annotations display characteristic aglycone peaks of triterpenoid saponins that have not been reported before in *C. sativa, C. hirsuta*, and *C. rubella*.

### Root and leaf tissue show distinct degrees of conservation in metabolic fingerprints

To investigate tissue-specific conservation and composition across supertribes and species, compound class annotations were assessed in leaf and root tissue profiles across supertribes and species. Positive and negative ionization modes revealed distinct patterns of specialized metabolite class distribution in both tissue types. In positive ionization mode, 2,206 features were identified in leaf tissue and 1,675 features in root tissue across all 14 species, with 1,237 features shared between both tissues (Fig. 7a). In negative ionization mode, 1,878 metabolite features were leaf-specific, and 1,327 were root-specific, while 902 were extracted from both tissues (Fig. 7b). The largest number of unique metabolite features exclusively obtained from root tissue was found in *B. vulgaris* in negative ionization mode (207, 59.7%, Fig S2). Notably, the root tissue profile of *B. vulgaris* exhibited a higher proportion of unique metabolite features than in leaf tissue in both ionization modes, whereas most other species showed the opposite trend (Fig. 7). The shared metabolite profile across all species displays a comparable composition of compound classes across tissues and ionization modes, representing 4.2% of all detected features in leaf and in root under positive mode, and 2.2% in leaf and 2.6% in root under negative mode (Fig. 7). Similarly, on the level of supertribes, tissue profiles shared between Camelinodae, Brassicodae and outgroup are comparable in positive (535 features, 15.4% in leaf and 477 features, 16.4% in root) and negative (285 features, 10.2% in leaf, 288 features, 13.0% in root) ionization modes. Adding the dimension of tissue revealed distinct degrees of conservation in species. The largest number (68, 5.4%) of metabolite features shared by two species was found in the Camelinodae (*C. sativa* and *C. rubella*) in root tissue under positive ionization mode, dominated by alkaloid and terpenoid annotations. In contrast, the shared metabolite features between *B. carinata, B. nigra* and *B. oleracea* belong to the leaf tissue profiles under negative ionization mode, dominated by amino acids, peptides, shikimates and phenylpropanoids (Fig. 7b1)

**Figure 7.**
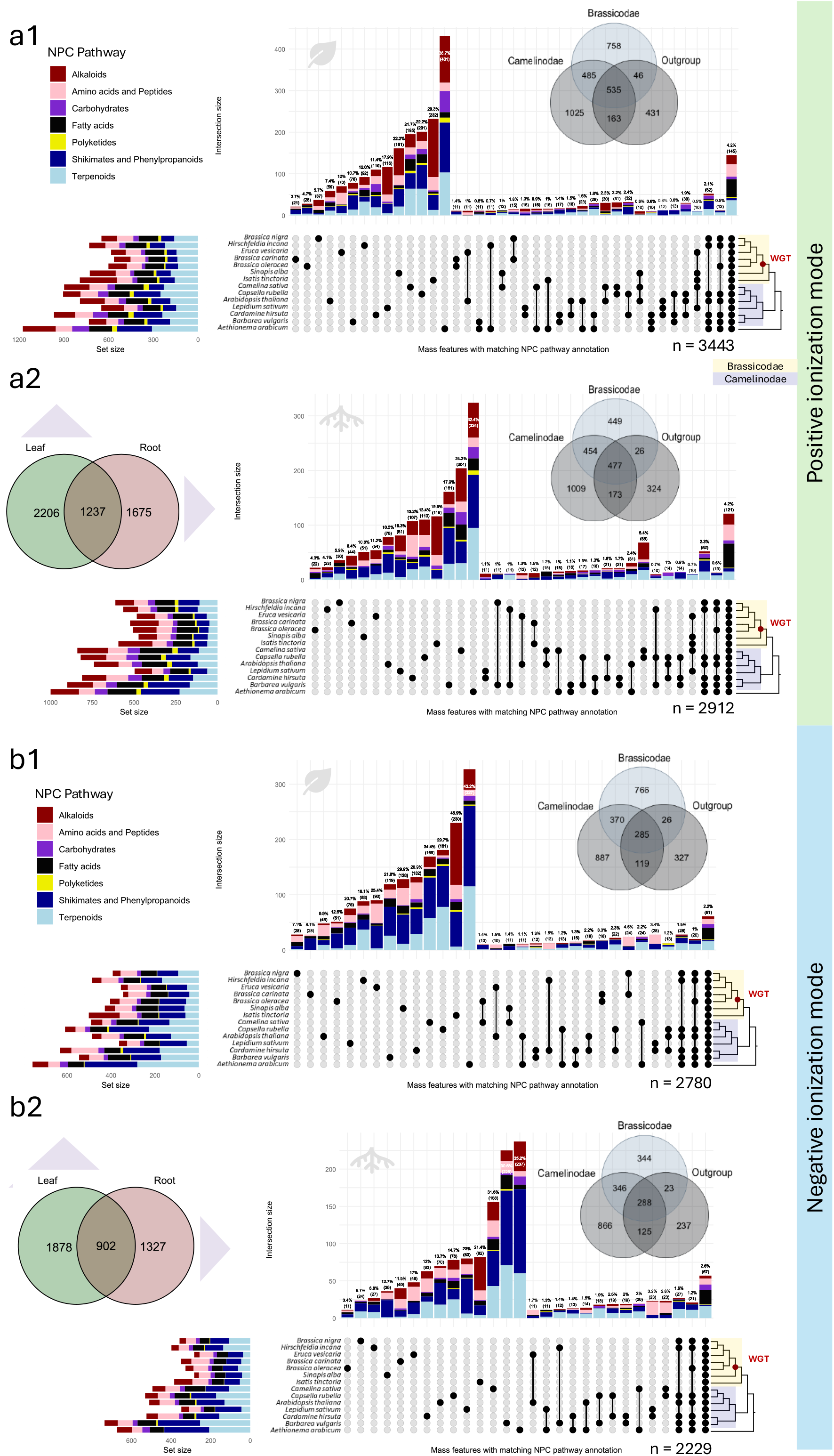
Distribution of mass features across leaf and root tissue in 14 Brassicaceae species.Untargeted metabolomics (LC-MS/MS) data was filtered for consistent annotations on the NP.Classifier “pathway” chemical ontology level. Red dots on phylogenetic trees indicate the Whole Genome Triplication (WGT) event in the Brassicodae clade. Venn diagrams show overlaps of aligned features across supertribes. **(a)** Metabolite features annotated in positive ionization mode in (**a1**) leaf and (**a2**) root tissue. **(b)** Metabolite features annotated in negative ionization mode in (**b1**) leaf and (**b2**) root tissue.

Our results indicate tissue-specific conservation of metabolic fingerprints in related plant species, underscoring the importance of spatial variation, and its implications for gene regulation in plant specialized metabolite biosynthesis. However, we suggest to interpret observed variation in leaf and root tissue metabolomes in the context of plant ontogeny, as recently highlighted for *Solanum dulcamara* (Anaia *et al*., 2025). Hence, metabolic fingerprinting may reveal distinct patterns of phytochemical relatedness across tissues depending on plant developmental stages.

### Chemical taxonomy across 17 species aligns largely with species phylogeny when combining leaf and root tissue profiles

LC-MS/MS data of leaf and root tissue extracts from *Rorippa islandica, Brassica juncea*, and *Arabis alpina* were integrated as described below with spectra obtained from tissues of 14 species listed in Table 1. *A. alpina* represents an additional outgroup to the Camelinodae supertribe, while *R. islandica* and *B. juncea* were added to the Camelinodae and Brassicodae supertribes, respectively (Fig. 8a). Batches of LC-MS/MS data were aligned using the MS-Cluster algorithm within the classical molecular networking workflow in GNPS2. MS-Cluster computes consensus spectra, disregarding retention time shifts across batches. The resulting classical molecular networks of 17 species across leaf and root tissue consist of 14,329 nodes in positive, and 11,271 nodes in negative ionization mode. Filtering for high-confidence annotations computed with SIRIUS v6.0.1 (cutoff: Confidence Exact Score ≥0.6) resulted in 1,048 compounds combining positive and negative ionization mode, and across leaf and root tissues of 17 species listed in Table 1.

**Figure 8.**
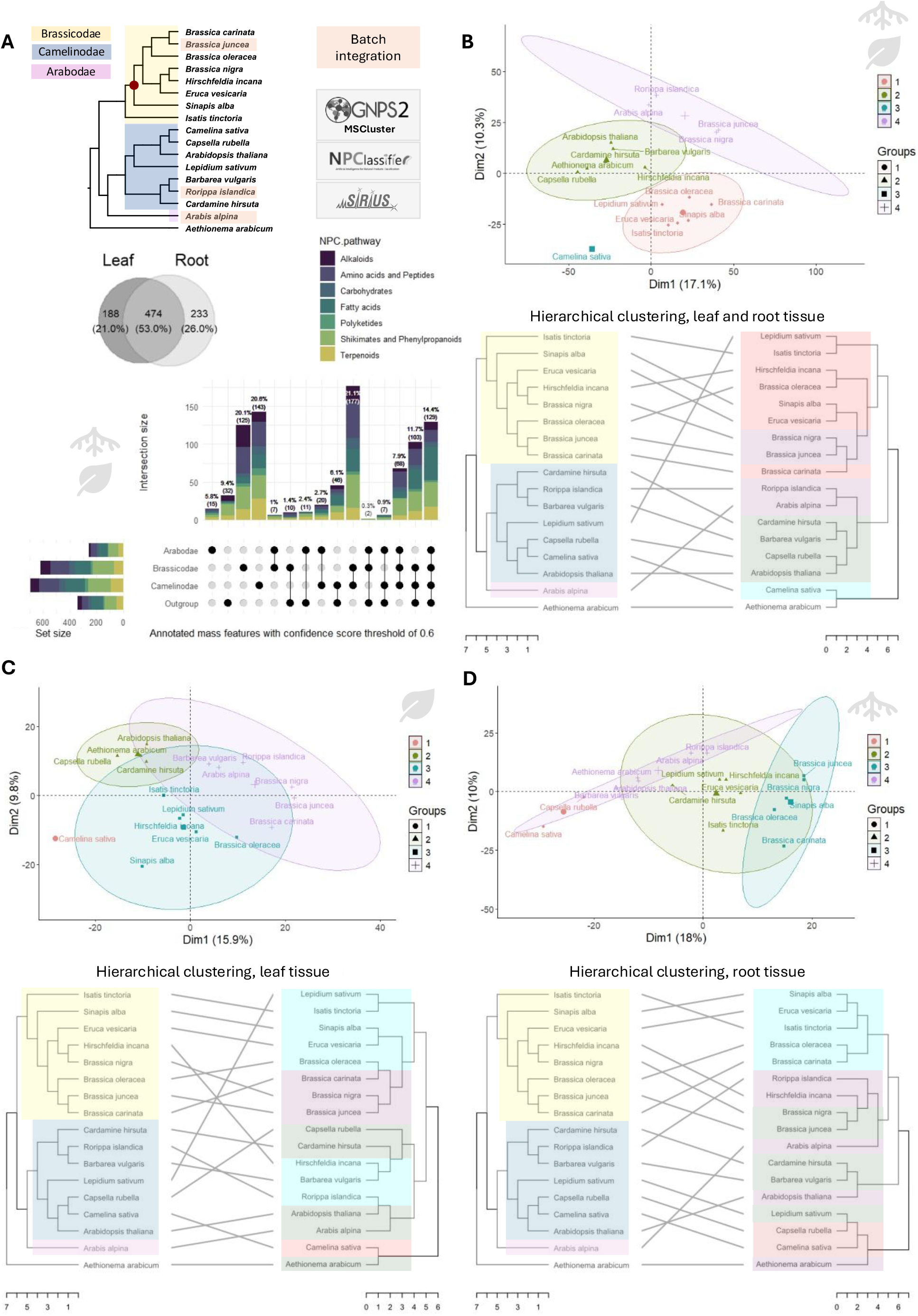
Hierarchical clustering of structural annotations in 17 species across leaf and root tissue and across two batches of LC-MS/MS data. Untargeted LC-MS/MS data was aligned by applying the MS-Cluster algorithm on GNPS2, and subsequently filtering for structural annotations obtained in SIRIUS with a Confidence Exact Score threshold of 0.6. Multiple identical structural annotations potentially resulting from in-source fragmentation and adduct formation were grouped. **A:** Integration of batches including 14 species (batch “a”) and three species (batch “b”). Venn diagram of shared and unique annotations in leaf and root tissue, and upset plot showing the distribution across NP.Classifier “pathway” annotations. **B:** PCA and hierarchical clustering of features found across leaf and root tissue. **C:** PCA and hierarchical clustering of features found in leaf tissue. **D:** PCA and hierarchical clustering of features found in root tissue.

Compound structure annotations with confidence exact score ≥0.6 obtained in SIRIUS for consensus spectra of mass features were then combined, to merge similar annotations (by matching InChI) across multiple features, resulting in 895 unique structure annotations in a precursor m/z range of 130.050 to 1337.720 Da (Material S5). The total number of unique annotations in root tissue (n = 233, 26.0%) was comparable with the number of features annotated in leaf tissue (188, 21.0%). Additionally, the largest proportion of annotated metabolite features is shared across tissues between Camelinodae and Brassicodae supertribes (177, 21.1%, Fig. 8a). Hierarchical complete clustering of the resulting set of 895 features was followed by the construction of tanglegrams for comparison with species phylogeny according to Hendriks *et al*. (2024) using mass features from both leaf and root tissues (Fig. 8b). Furthermore, hierarchical complete clustering and resulting tanglegrams were compared for the total number of features extracted from leaf (n = 662) (Fig. 8c) and root tissue (n = 707) (Fig. 8d), including features extracted from both tissue types (n = 474). The numbers of annotated metabolite features extracted exclusively from leaf (188) and root (233) tissue are given in Table 2 and in the Venn diagram shown in Fig 8a.

**Table 2.** Number of metabolite features obtained for integrated batches including LC-MS/MS data of leaf and root extracts of 17 species.

|  |  |
| --- | --- |
| **Unfiltered consensus spectra** | 11,271 in positive mode<br>14,329 in negative mode |
| **InChI with confidence exact score >0.6 obtained in SIRIUS v.6.1.0** | 1,048 |
| **Unique InChI in consensus spectra** |  |
| Leaf-specific profile | 188 |
| **Leaf profile (total)** | $188 + 474 = 662$ |
| Root-specific profile | 233 |
| **Root profile (total)** | $233 + 474 = 707$ |
| Shared profile (leaf/root) | 474 |
| **Total (all features)** | 895 |

Clustering of all features, across both tissue types, resulted in four clusters in the PCA, explaining 27.4% of variance in the first two dimensions. Notably, the outgroup is represented by *C. sativa*, while *A. arabicum* clusters with species from the Camelinodae supertribe in cluster 2. Species belonging to the Brassicodae supertribe cluster in two distinct groups, together with *A. alpina* and *Lepidium sativum* in cluster 1 and 4, respectively. The corresponding tanglegram largely aligns with species phylogeny for the complete filtered dataset, including both tissue types, while *L. sativum* and *A. alpina* were found in distinct clusters (Fig. 8b).

Tissue-specific hierarchical clustering of features revealed a similar displacement of *L. sativum* and *A. alpina* for leaf and root tissue. Clustering restricted to features found in leaf tissue reveals separation of *Sinapis alba* in cluster 3 from *Brassica* species in cluster 4, primarily along the first dimension explaining 15.9% of variance (Fig. 8C). Notably, clustering of annotated features found in root tissue indicates proximity of *C. sativa* and *C. rubella* in the same cluster, representing the outgroup to the otherwise largely preserved Camelinodae and Brassicodae supertribes (Fig. 8d). Although clusters for tissue-specific datasets largely overlap, *C. sativa* persistently appears as an outgroup in all three clustering scenarios. In addition, the consistent displacement of *L. sativum* clustering with *I. tinctoria* indicates phytochemical relatedness. Incorporating distinct species in the genus *Lepidium* and *Isatis*, and wild relatives next to domesticated accessions may further resolve tissue-specific chemotaxonomic patterns within the Lepidae tribe.

Our results on comparative chemotaxonomy across two batches of untargeted LCMS/MS data show that phylogenetic distance can be largely recovered from integrated data across batches, tissues, and ionization modes. Although structure annotation, and hence comparative analysis of compound class distribution is severely limited for this type of analysis, metabolic distance based on hierarchical clustering reflected phylogenetic distance to a large extent. Furthermore, tissue-specific clustering highlighted potentially divergent conservation of tissue-specific profiles. Discrete evolutionary pressures on plant tissues have been suspected earlier to result in conserved metabolite profiles incongruent with species phylogeny (Nguyen *et al*., 2025). When profiles of leaf and root tissue were combined, clustering of Brassicodae and Camelinodae species in distinct groups was retained, with two exceptions, namely the placements of *L. sativum* and *C. sativa* in Brassicodae and outgroup, respectively. The extension of chemical taxonomy to other supertribes and different plant families may increase the resolution to investigate the relative distance of *C. sativa* to Camelinodae species.

### Clustering of Brassica species reflects relationships of shared sub-genomes in descendants of *B. nigra* and *B. oleracea*

To investigate the metabolic distance among species within the *Brassica* genus, we included *B. juncea*, which harbors the A and B genomes derived from its progenitor species *B. rapa* and *B. nigra* (*Mutarda nigra*). In leaf tissue-specific hierarchical clustering, diploid *B. nigra* and *B. oleracea* group with the polyploids *B. juncea* (an AB genome species) and *B. carinata* (a BC genome species), respectively (Fig. 8c). All included *Brassica* species formed a distinct, monophyletic clade based on leaf tissue profiles, indicating conserved tissue-specific chemistry within this genus. In contrast, *B. nigra* and *B. juncea* harboring the B genome were split from *B. oleracea* and *B. carinata* harboring the C genome in root tissue-specific clustering (Fig. 8d). Interestingly, the B-genome species *B. nigra* groups with its close relative *H. incana* when only including root-specific profiles. However, when leaf and root tissue profiles are jointly analyzed, *H. incana* clusters with the C-genome species *B. oleracea* in a separate cluster, while *B. nigra, B. juncea* and *B. carinata* form a monophyletic clade (Fig. 8b).Consistent grouping of *B. nigra* and *B. juncea* suggests that the metabolite profile of *B. juncea* is consistently characterized by the shared B genome across leaf and root tissue. In contrast, *B. carinata* grouping consistently with *B. oleracea* suggests that the metabolite profile in *B. carinata* is more determined by the C genome rather than the B genome, particularly in root tissue.

We included merely a filtered set of metabolite features in our analysis, restricted to high-confidence structure predictions in SIRIUS. However, clustering of *B. nigra, B. juncea, B. carinata*, and *B. oleracea* is observed across all clustering scenarios, including unfiltered aligned metabolite features (Figure S2). The inclusion of *B. rapa* and *B. napus* covering A and AC genomes in metabolic fingerprinting for chemical taxonomy may provide further insights into the effects of sub-genome dominance of A versus B and C genomes in future research.

### Outlook and perspectives

To systematically investigate the composition of shared and unique metabolic profiles across the Brassicaceae, we conducted untargeted LC-MS/MS covering 14 species across two major supertribes (Brassicodae and Camelinodae) and two tissues (leaf and root). We assessed taxonomic relationships reflected by conserved metabolite profiles across species and tissues. Our results suggest that comparative metabolic fingerprinting reveals potential hotspots of phytochemical innovation. Current estimates of the total number of unique compounds in the plant kingdom are extrapolated based on a limited number of plant species, while not accounting for phylogenetic distances. We therefore advocate designing studies that systematically profile species across phylogenetic levels to improve estimates of remaining chemical diversity. Balanced, clade-wide sampling can reveal shared biochemical traits within lineages and help correct biases toward model and crop species. Furthermore, we established a computational pipeline for pre-processing and annotation of positive and negative ionization mode LC-MS/MS data integrating *in silico* structure, substructure, and compound class annotation tools. We found that library matches and consensus predictions on chemical ontology levels overlap with predicted substructures (Mass2Motifs), highlighted for the triterpenoid compound class. Our results also align with previously described trends in phytochemical diversity across multiple species, as found for alkaloids and triterpenoid saponins in *I. tinctoria* and *B. vulgaris*, respectively.

Since specialized metabolites are often stress-induced, metabolic fingerprinting reflects only a snapshot of plant chemical diversity, underscoring the importance of systematically eliciting hormone-induced stress responses in experimental designs to further probe the plant biochemical repertoire. Furthermore, we note that our analytical approach does not provide unambiguous proof of structural identities; however, our framework was designed to combine inputs from multiple different sources to provide consensus annotations, making annotations on chemical ontology levels more reliable. Ultimately, our experimental and computational workflows provide a modular and extendible framework for systematic metabolic fingerprinting to study the evolution of plant natural product biosynthesis.

## Acknowledgements

This work was supported by the Netherlands Organization for Scientific Research (NWO) Vidi Grant VI.Vidi.213.183 to M.H.M. The authors would like to thank Ric de Vos and Bert Schipper for their help and comments on the experimental design and metabolite extraction protocols, and for establishing the analytical platforms. Further, the authors would like to thank Nam Hoang for providing valuable feedback on the determination of candidate species implemented in this research, and for his helping hands in preliminary experiments.

## Competing interests

The authors declare the following competing financial interest(s): M.H.M. is a member of the scientific advisory boards of Hexagon Bio and Hothouse Therapeutics. J.J.J.vdH. is member of the scientific advisory board of NAICONS Srl., Milano, Italy and consults for Corteva Agriscience, Indianapolis, IN, USA. All other authors declare no competing interests.

## Author contributions

F.W., M.H.M., K.B., and J.J.J.vdH. designed the study. F.W. and T.W. conducted the experiments and collected the data. F.W. and J.J.J.vdH. conducted LC-MS/MS data analysis. F.W. created figures and tables. F.W. wrote the initial draft of the manuscript, and all authors contributed substantially to revisions. No generative AI (LLMs) was used for writing this manuscript.

## Data availability

The LC-MS/MS data collected in this study has been submitted to MetaboLights (MTBLS14270) and MassIVE (MSV000101479). Scripts are available on Github (https://github.com/capsicumbaccatum/Metabolic_fingerprinting_consensus_annotation). Supplementary data, such as intermediate results can be found on Zenodo (DOI: 10.5281/zenodo.19391098).

## Supporting Information

A list of the Supplementary Files is provided here below.

### Supplementary_figure_S1-3. pdf

**Fig**. **S1** Total number and overlap of computational annotation tools (SIRIUS, MS2Query and Dreams against the MassSpecGym library) in positive and negative ionization mode

**Fig. S2** Tanglegrams showing hierarchical clustering of aligned filtered and unfiltered metabolite features across 17 species

**Fig. S3** Upset plots showing shared and unique annotated metabolite features exclusively found in leaf and root tissue of 14 species

### Supplementary material available on Zenodo

**Material S1:** Instrument settings and mass exclusion list for LC-MS/MS in positive and negative ionization mode

**(DOI: 10.5281/zenodo.19391098)**

**Material S2:** mzmine 4.8.5 batchfile in XML format, including all settings for feature extraction and alignment, and aligned feature-lists

**Material S3:** MS2LDA analysis: config file and full results in positive and negative ionization mode

**Material S4:** HTML summaries of computed consensus annotations across SIRIUS v6.1.0, MS2Query v1.0 and Dreams v1.0 against the MassSpecGym library

**Material S5:** Raw and filtered aligned mass features across two batches, including 17 species (leaf and root), using MSCluster. All intermediate files are provided

**Material S6 :** Complete annotation table, including all annotation levels (SIRIUS, MS2Query and Dreams against the MassSpecGym library, GNPS library matches, Mass2Motifs, total abundance per sample, taxonomic levels, feature Id) and fully annotated GNPS2 FBMN

## Notes

https://doi.org/10.5281/zenodo.19391098

https://github.com/capsicumbaccatum/Metabolic_fingerprinting_consensus_annotation

