## Supplementary figure S1-3 for "Systematic mass-spectrometry-guided metabolic fingerprinting elucidates diversity of specialized metabolites across the Brassicaceae"

**Positive ionization mode**

(a)

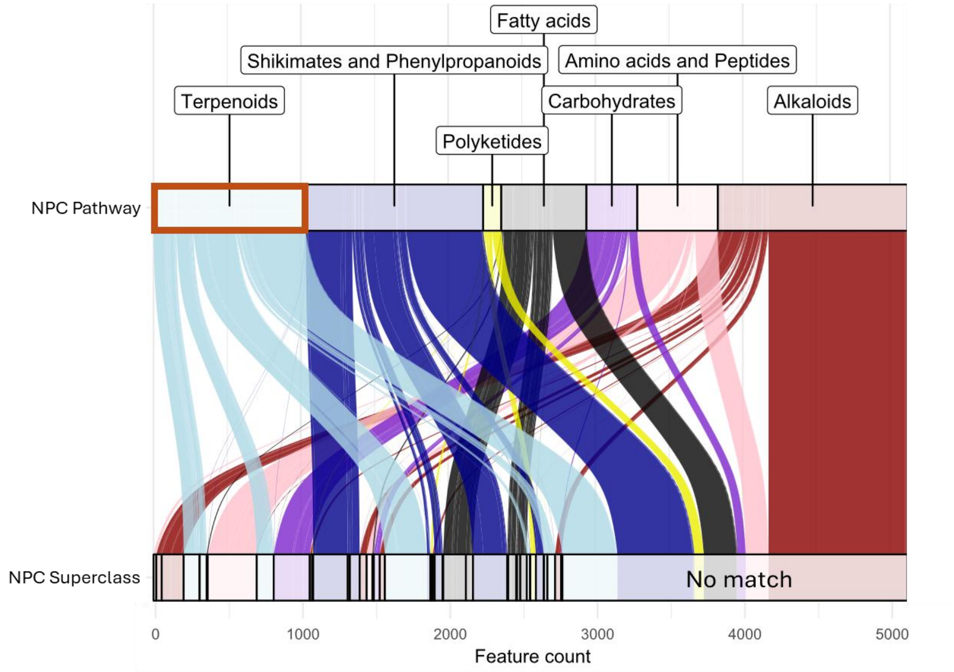


**Consensus annotation**

- Pathway (all) 5,118
- Pathway: Terpenoids 1,032
- Superclass: Triterpenoids 108

**Negative ionization mode**

(b)

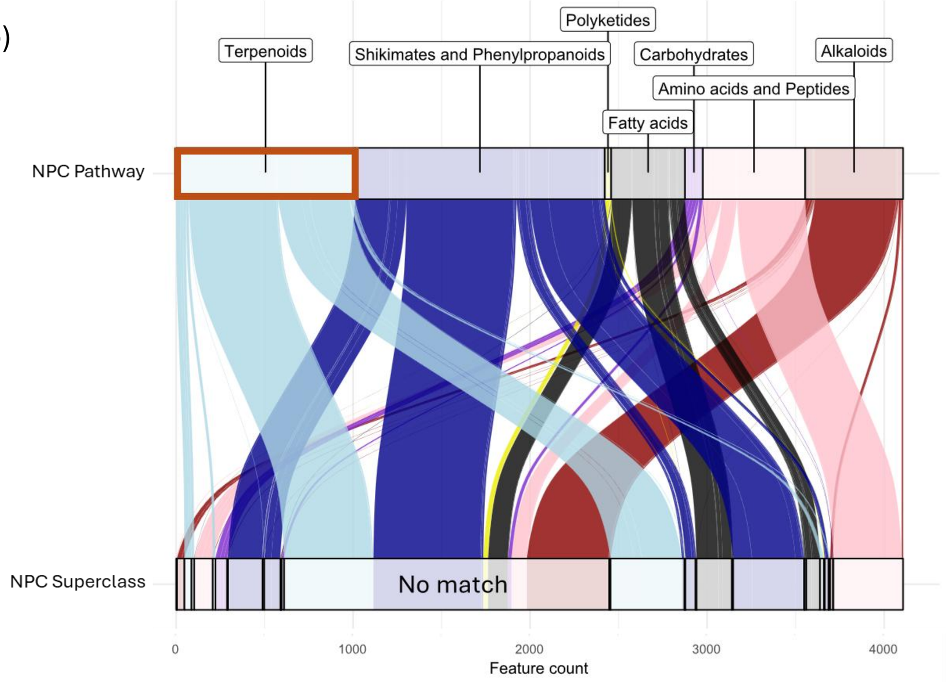


**Consensus annotation**

- Pathway (all) 4,107
- Pathway: Terpenoids 1,011
- Superclass: Triterpenoids 40

**Fig. S1** Total number and overlap of computational annotation tools (SIRIUS v6.1.0, MS2Query v1.0 and Dreams v1.0 against the MassSpecGym library) in (a) positive and (b) negative ionization mode across 14 species and two tissues (leaf and root).

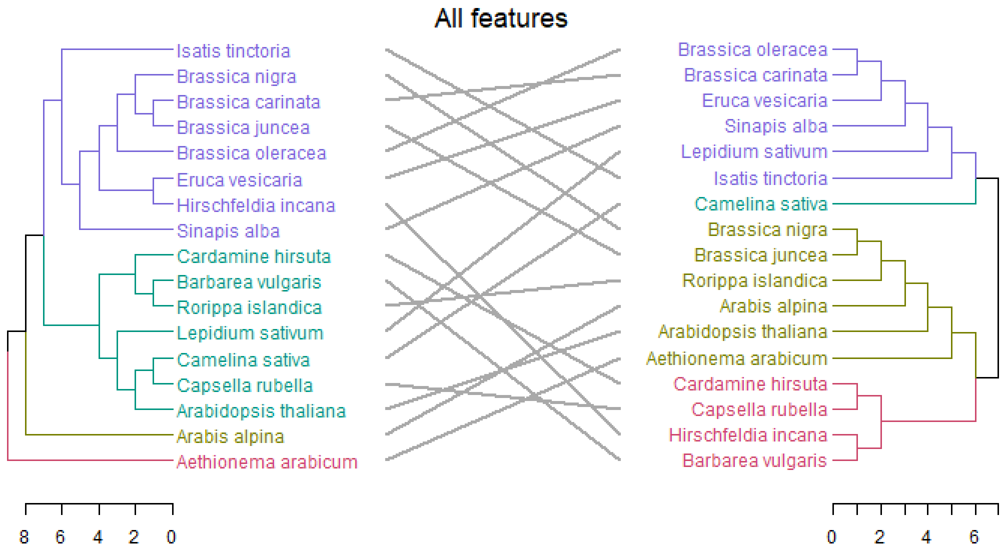
All features
Isatis tinctoria
Brassica nigra
Brassica carinata
Brassica juncea
Brassica oleracea
Eruca vesicaria
Hirschfeldia incana
Sinapis alba
Cardamine hirsuta
Barbarea vulgaris
Rorippa islandica
Lepidium sativum
Camelina sativa
Capsella rubella
Arabidopsis thaliana
Arabis alpina
Aethionema arabicum
8 6 4 2 0
Brassica oleracea
Brassica carinata
Eruca vesicaria
Sinapis alba
Lepidium sativum
Isatis tinctoria
Camelina sativa
Brassica nigra
Brassica juncea
Rorippa islandica
Arabis alpina
Arabidopsis thaliana
Aethionema arabicum
Cardamine hirsuta
Capsella rubella
Hirschfeldia incana
Barbarea vulgaris
0 2 4 6

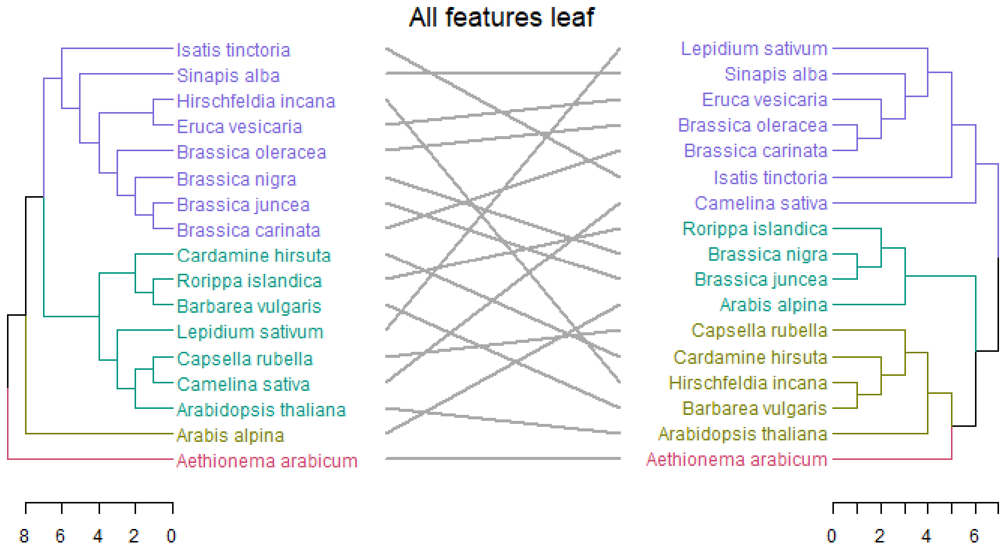
All features leaf
Isatis tinctoria
Sinapis alba
Hirschfeldia incana
Eruca vesicaria
Brassica oleracea
Brassica nigra
Brassica juncea
Brassica carinata
Cardamine hirsuta
Rorippa islandica
Barbarea vulgaris
Lepidium sativum
Capsella rubella
Camelina sativa
Arabidopsis thaliana
Arabis alpina
Aethionema arabicum
8 6 4 2 0
Lepidium sativum
Sinapis alba
Eruca vesicaria
Brassica oleracea
Brassica carinata
Isatis tinctoria
Camelina sativa
Rorippa islandica
Brassica nigra
Brassica juncea
Arabis alpina
Capsella rubella
Cardamine hirsuta
Hirschfeldia incana
Barbarea vulgaris
Arabidopsis thaliana
Aethionema arabicum
0 2 4 6

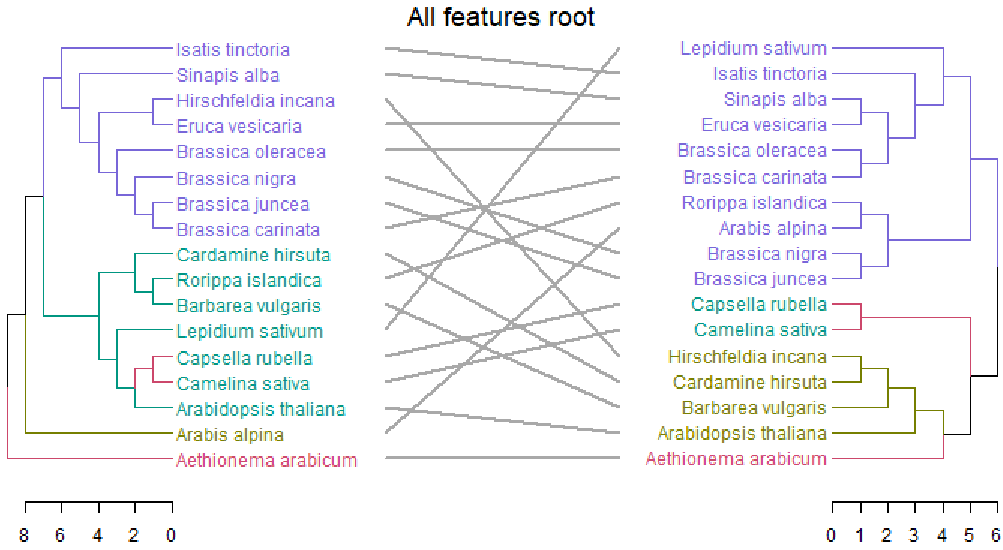
All features root
Isatis tinctoria
Sinapis alba
Hirschfeldia incana
Eruca vesicaria
Brassica oleracea
Brassica nigra
Brassica juncea
Brassica carinata
Cardamine hirsuta
Rorippa islandica
Barbarea vulgaris
Lepidium sativum
Capsella rubella
Camelina sativa
Arabidopsis thaliana
Arabis alpina
Aethionema arabicum
8 6 4 2 0
Lepidium sativum
Isatis tinctoria
Sinapis alba
Eruca vesicaria
Brassica oleracea
Brassica carinata
Rorippa islandica
Arabis alpina
Brassica nigra
Brassica juncea
Capsella rubella
Camelina sativa
Hirschfeldia incana
Cardamine hirsuta
Barbarea vulgaris
Arabidopsis thaliana
Aethionema arabicum
0 1 2 3 4 5 6

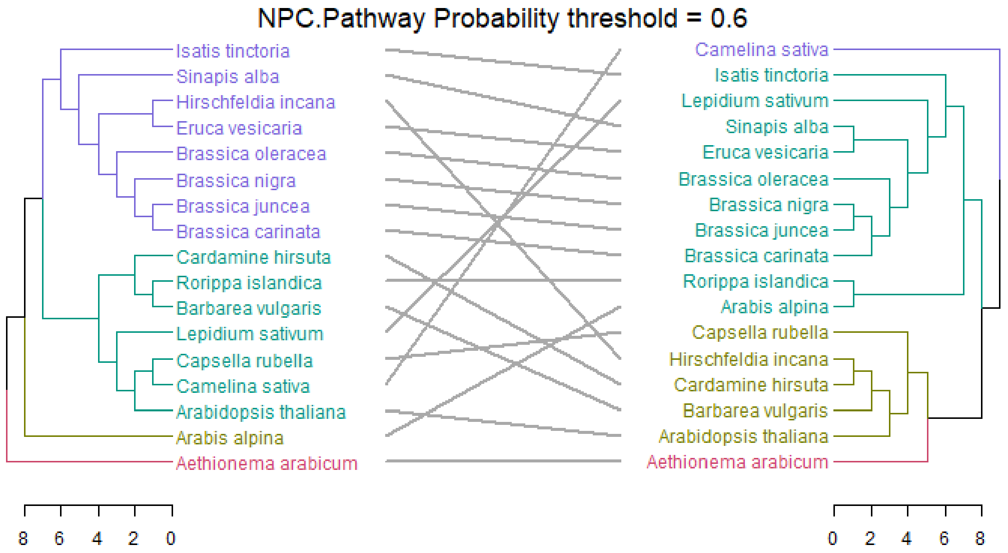
NPC.Pathway Probability threshold = 0.6
Isatis tinctoria
Sinapis alba
Hirschfeldia incana
Eruca vesicaria
Brassica oleracea
Brassica nigra
Brassica juncea
Brassica carinata
Cardamine hirsuta
Rorippa islandica
Barbarea vulgaris
Lepidium sativum
Capsella rubella
Camelina sativa
Arabidopsis thaliana
Arabis alpina
Aethionema arabicum
Camelina sativa
Isatis tinctoria
Lepidium sativum
Sinapis alba
Eruca vesicaria
Brassica oleracea
Brassica nigra
Brassica juncea
Brassica carinata
Rorippa islandica
Arabis alpina
Capsella rubella
Hirschfeldia incana
Cardamine hirsuta
Barbarea vulgaris
Arabidopsis thaliana
Aethionema arabicum
8 6 4 2 0
0 2 4 6 8

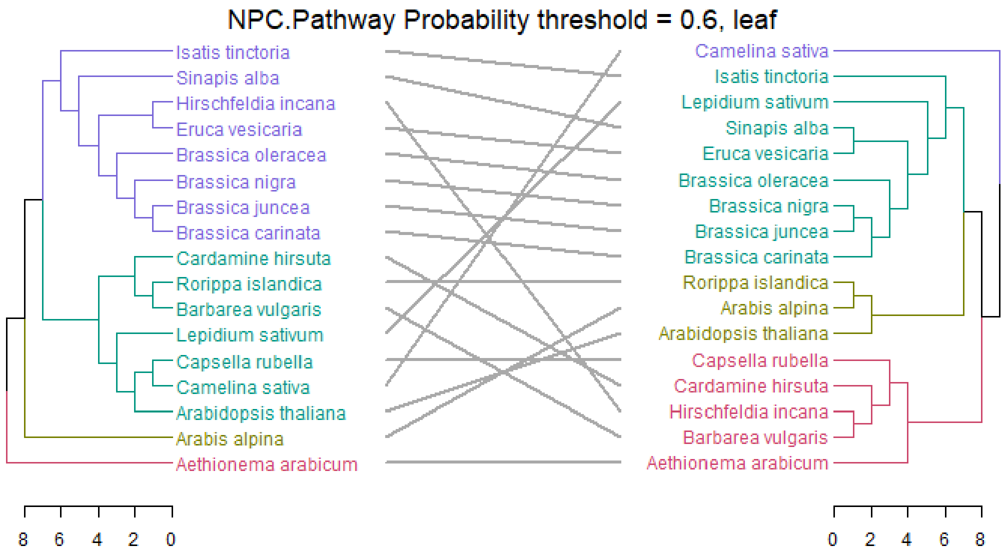
NPC.Pathway Probability threshold = 0.6, leaf
Isatis tinctoria
Sinapis alba
Hirschfeldia incana
Eruca vesicaria
Brassica oleracea
Brassica nigra
Brassica juncea
Brassica carinata
Cardamine hirsuta
Rorippa islandica
Barbarea vulgaris
Lepidium sativum
Capsella rubella
Camelina sativa
Arabidopsis thaliana
Arabis alpina
Aethionema arabicum
Camelina sativa
Isatis tinctoria
Lepidium sativum
Sinapis alba
Eruca vesicaria
Brassica oleracea
Brassica nigra
Brassica juncea
Brassica carinata
Rorippa islandica
Arabis alpina
Arabidopsis thaliana
Capsella rubella
Cardamine hirsuta
Hirschfeldia incana
Barbarea vulgaris
Aethionema arabicum
8 6 4 2 0
0 2 4 6 8

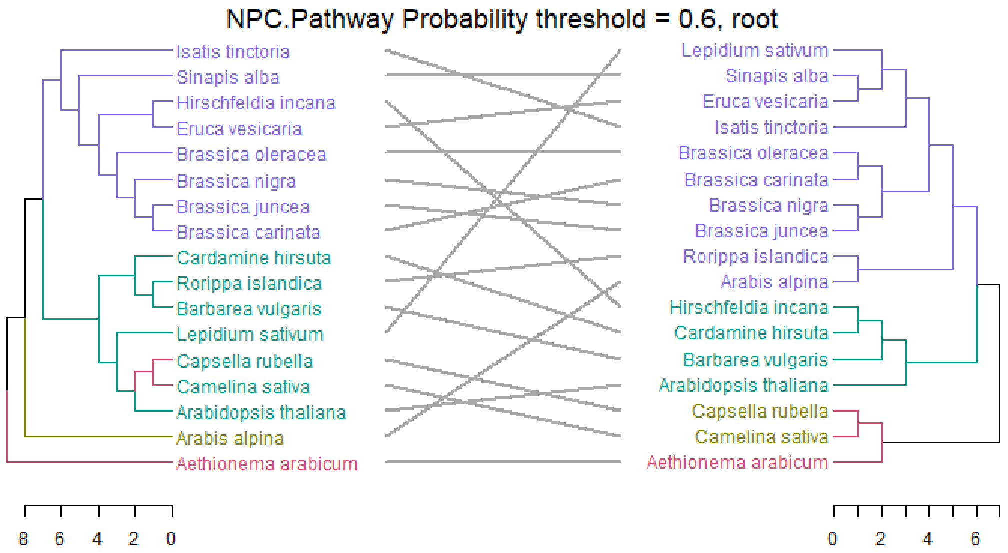
NPC.Pathway Probability threshold = 0.6, root
Isatis tinctoria
Sinapis alba
Hirschfeldia incana
Eruca vesicaria
Brassica oleracea
Brassica nigra
Brassica juncea
Brassica carinata
Cardamine hirsuta
Rorippa islandica
Barbarea vulgaris
Lepidium sativum
Capsella rubella
Camelina sativa
Arabidopsis thaliana
Arabis alpina
Aethionema arabicum
Lepidium sativum
Sinapis alba
Eruca vesicaria
Isatis tinctoria
Brassica oleracea
Brassica carinata
Brassica nigra
Brassica juncea
Rorippa islandica
Arabis alpina
Hirschfeldia incana
Cardamine hirsuta
Barbarea vulgaris
Arabidopsis thaliana
Capsella rubella
Camelina sativa
Aethionema arabicum
8 6 4 2 0
0 2 4 6

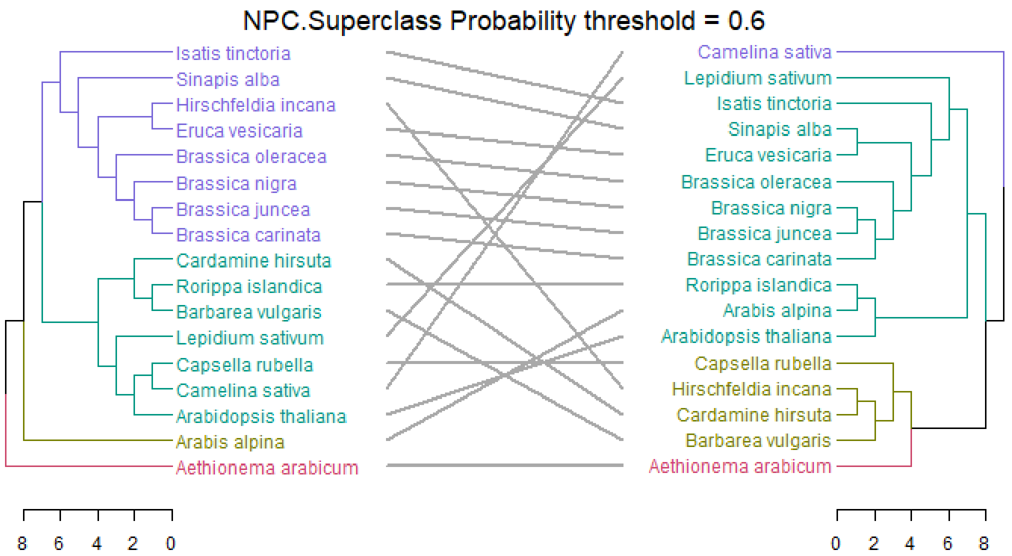
NPC.Superclass Probability threshold = 0.6
Isatis tinctoria
Sinapis alba
Hirschfeldia incana
Eruca vesicaria
Brassica oleracea
Brassica nigra
Brassica juncea
Brassica carinata
Cardamine hirsuta
Rorippa islandica
Barbarea vulgaris
Lepidium sativum
Capsella rubella
Camelina sativa
Arabidopsis thaliana
Arabis alpina
Aethionema arabicum
Camelina sativa
Lepidium sativum
Isatis tinctoria
Sinapis alba
Eruca vesicaria
Brassica oleracea
Brassica nigra
Brassica juncea
Brassica carinata
Rorippa islandica
Arabis alpina
Arabidopsis thaliana
Capsella rubella
Hirschfeldia incana
Cardamine hirsuta
Barbarea vulgaris
Aethionema arabicum
8 6 4 2 0
0 2 4 6 8

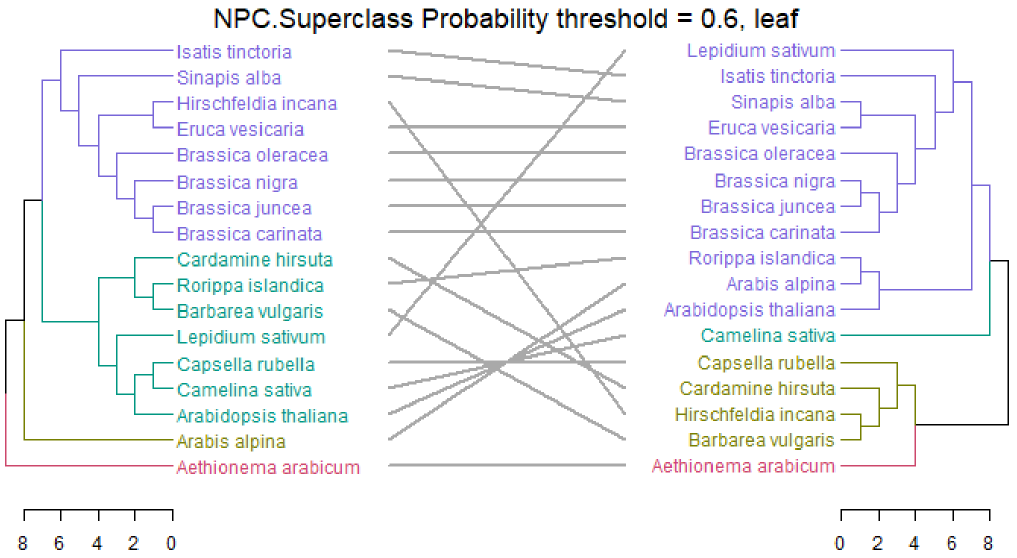
NPC.Superclass Probability threshold = 0.6, leaf
Isatis tinctoria
Sinapis alba
Hirschfeldia incana
Eruca vesicaria
Brassica oleracea
Brassica nigra
Brassica juncea
Brassica carinata
Cardamine hirsuta
Rorippa islandica
Barbarea vulgaris
Lepidium sativum
Capsella rubella
Camelina sativa
Arabidopsis thaliana
Arabis alpina
Aethionema arabicum
Lepidium sativum
Isatis tinctoria
Sinapis alba
Eruca vesicaria
Brassica oleracea
Brassica nigra
Brassica juncea
Brassica carinata
Rorippa islandica
Arabis alpina
Arabidopsis thaliana
Camelina sativa
Capsella rubella
Cardamine hirsuta
Hirschfeldia incana
Barbarea vulgaris
Aethionema arabicum
8 6 4 2 0
0 2 4 6 8

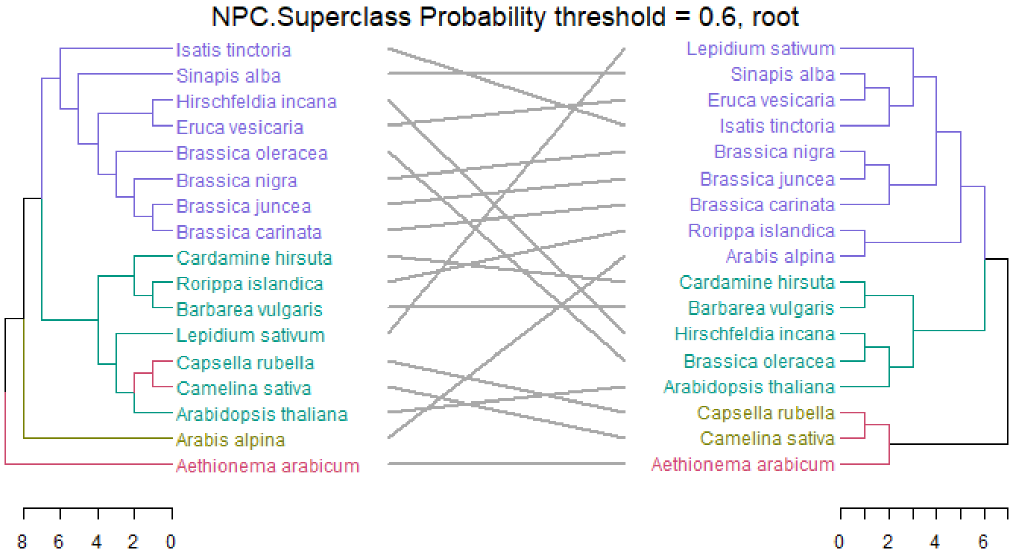
NPC.Superclass Probability threshold = 0.6, root
Isatis tinctoria
Sinapis alba
Hirschfeldia incana
Eruca vesicaria
Brassica oleracea
Brassica nigra
Brassica juncea
Brassica carinata
Cardamine hirsuta
Rorippa islandica
Barbarea vulgaris
Lepidium sativum
Capsella rubella
Camelina sativa
Arabidopsis thaliana
Arabis alpina
Aethionema arabicum
Lepidium sativum
Sinapis alba
Eruca vesicaria
Isatis tinctoria
Brassica nigra
Brassica juncea
Brassica carinata
Rorippa islandica
Arabis alpina
Cardamine hirsuta
Barbarea vulgaris
Hirschfeldia incana
Brassica oleracea
Arabidopsis thaliana
Capsella rubella
Camelina sativa
Aethionema arabicum
8 6 4 2 0
0 2 4 6

NPC.Class Probability threshold = 0.6

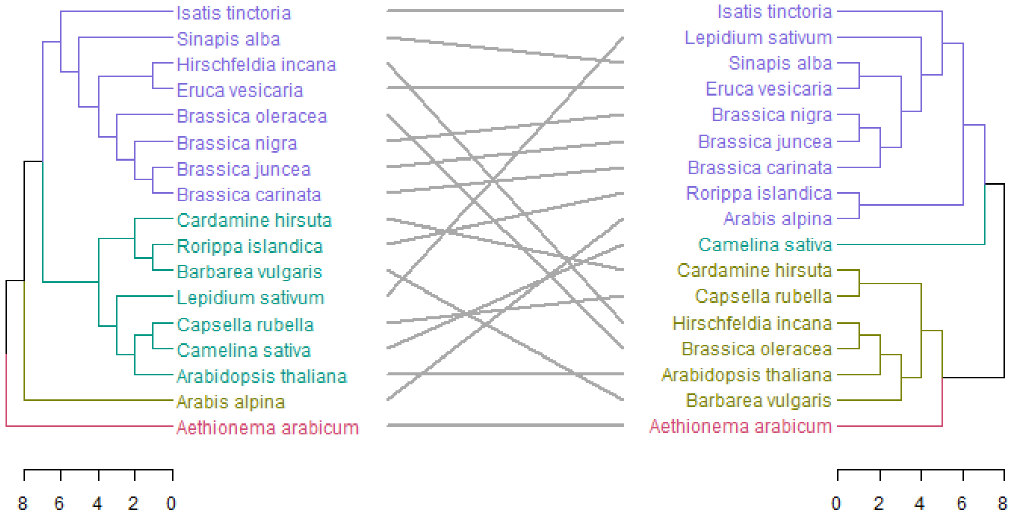


NPC.Class Probability threshold = 0.6, leaf

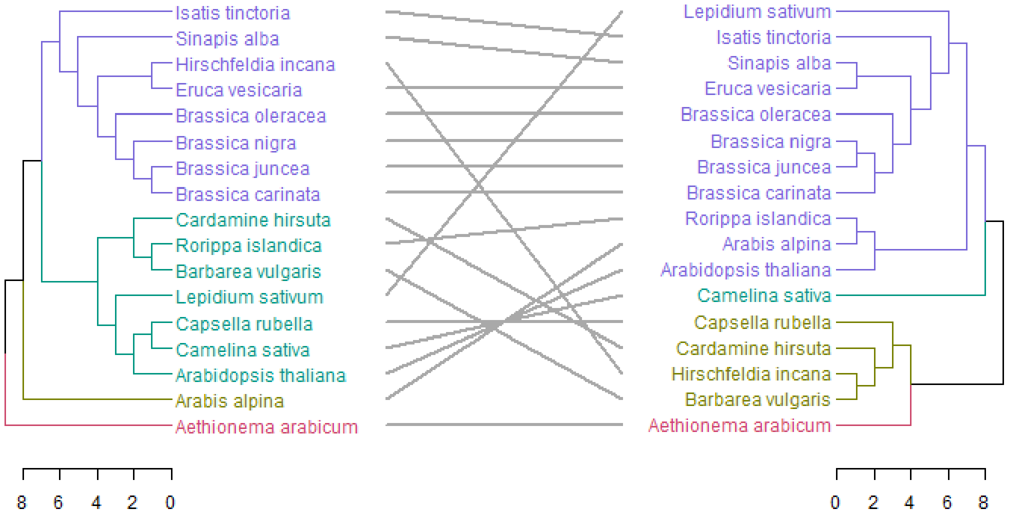


NPC.ClassProbability threshold = 0.6, root

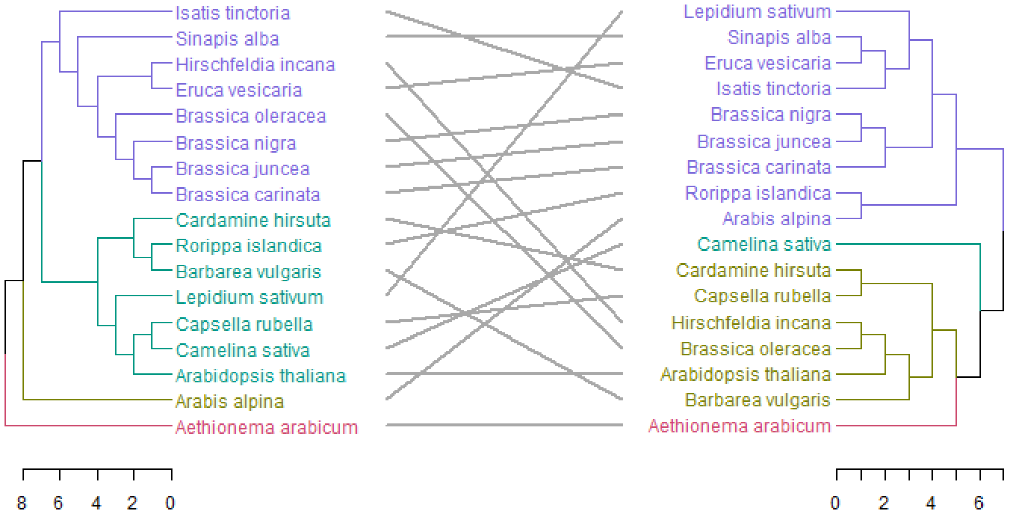

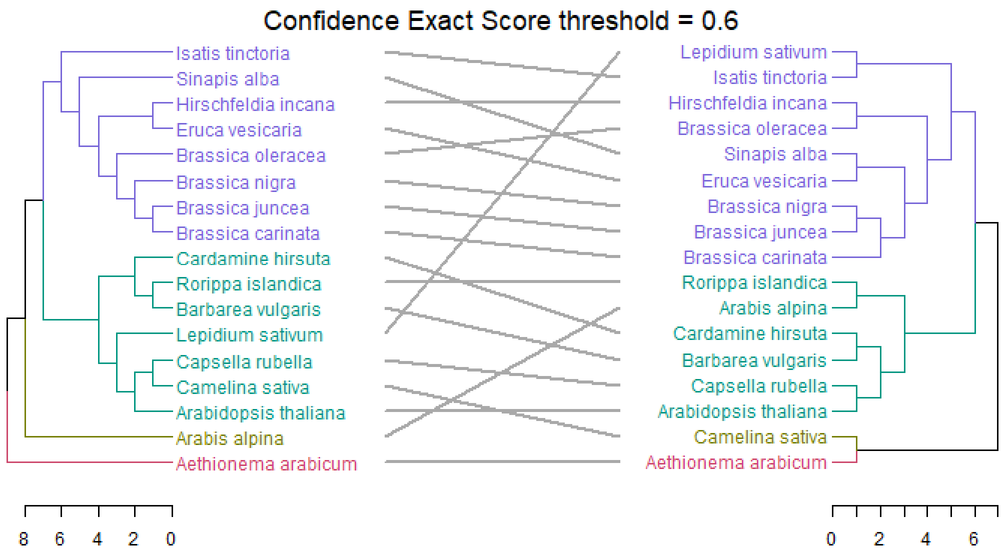
Confidence Exact Score threshold = 0.6
Isatis tinctoria
Sinapis alba
Hirschfeldia incana
Eruca vesicaria
Brassica oleracea
Brassica nigra
Brassica juncea
Brassica carinata
Cardamine hirsuta
Rorippa islandica
Barbarea vulgaris
Lepidium sativum
Capsella rubella
Camelina sativa
Arabidopsis thaliana
Arabis alpina
Aethionema arabicum
Lepidium sativum
Isatis tinctoria
Hirschfeldia incana
Brassica oleracea
Sinapis alba
Eruca vesicaria
Brassica nigra
Brassica juncea
Brassica carinata
Rorippa islandica
Arabis alpina
Cardamine hirsuta
Barbarea vulgaris
Capsella rubella
Arabidopsis thaliana
Camelina sativa
Aethionema arabicum
8 6 4 2 0
0 2 4 6

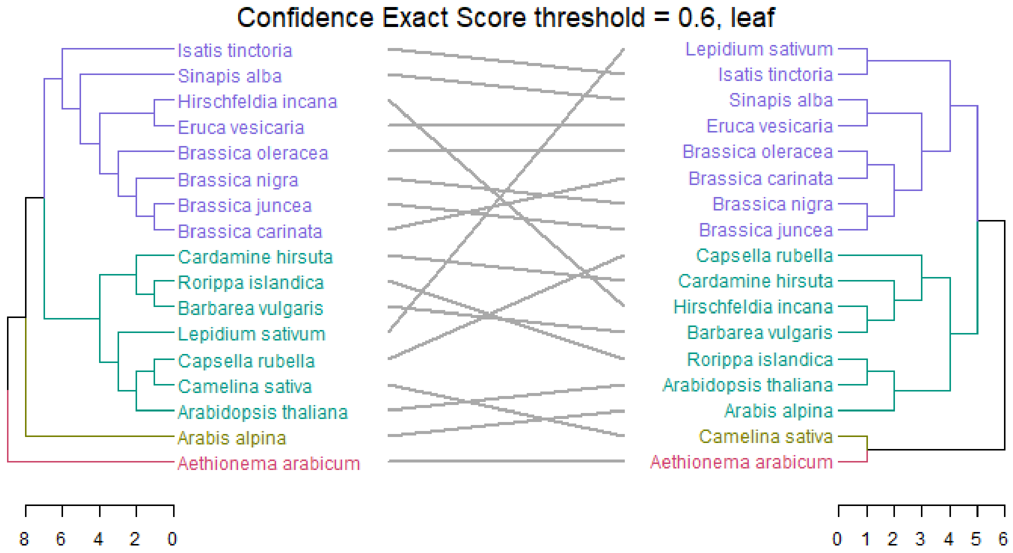
Confidence Exact Score threshold = 0.6, leaf
Isatis tinctoria
Sinapis alba
Hirschfeldia incana
Eruca vesicaria
Brassica oleracea
Brassica nigra
Brassica juncea
Brassica carinata
Cardamine hirsuta
Rorippa islandica
Barbarea vulgaris
Lepidium sativum
Capsella rubella
Camelina sativa
Arabidopsis thaliana
Arabis alpina
Aethionema arabicum
Lepidium sativum
Isatis tinctoria
Sinapis alba
Eruca vesicaria
Brassica oleracea
Brassica carinata
Brassica nigra
Brassica juncea
Capsella rubella
Cardamine hirsuta
Hirschfeldia incana
Barbarea vulgaris
Rorippa islandica
Arabidopsis thaliana
Arabis alpina
Camelina sativa
Aethionema arabicum
8 6 4 2 0
0 1 2 3 4 5 6

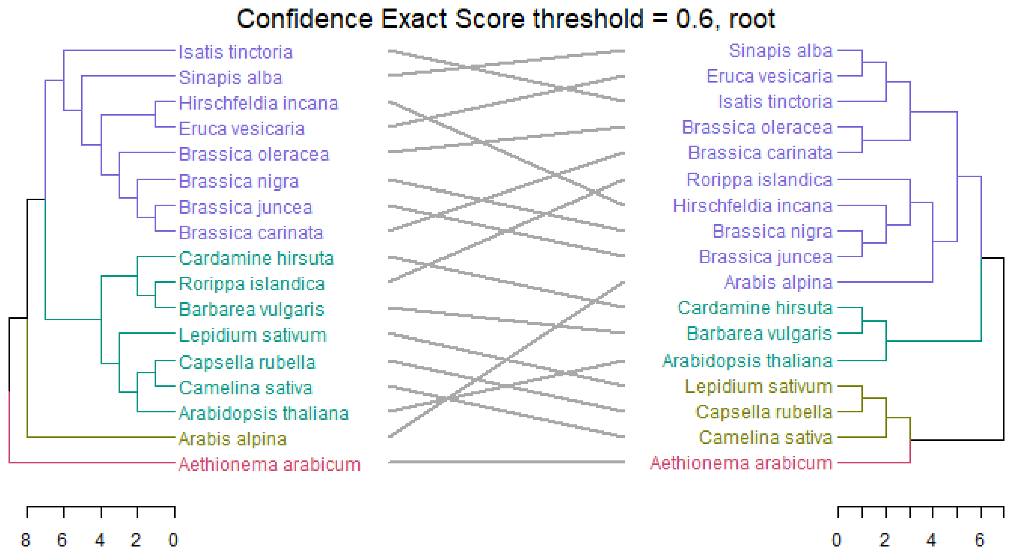
Confidence Exact Score threshold = 0.6, root
Isatis tinctoria
Sinapis alba
Hirschfeldia incana
Eruca vesicaria
Brassica oleracea
Brassica nigra
Brassica juncea
Brassica carinata
Cardamine hirsuta
Rorippa islandica
Barbarea vulgaris
Lepidium sativum
Capsella rubella
Camelina sativa
Arabidopsis thaliana
Arabis alpina
Aethionema arabicum
Sinapis alba
Eruca vesicaria
Isatis tinctoria
Brassica oleracea
Brassica carinata
Rorippa islandica
Hirschfeldia incana
Brassica nigra
Brassica juncea
Arabis alpina
Cardamine hirsuta
Barbarea vulgaris
Arabidopsis thaliana
Lepidium sativum
Capsella rubella
Camelina sativa
Aethionema arabicum
8 6 4 2 0
0 2 4 6

**Fig. S2** Tanglegrams showing hierarchical clustering of aligned filtered and unfiltered metabolite features across 17 species.

Positive ionization mode, leaf-specific:

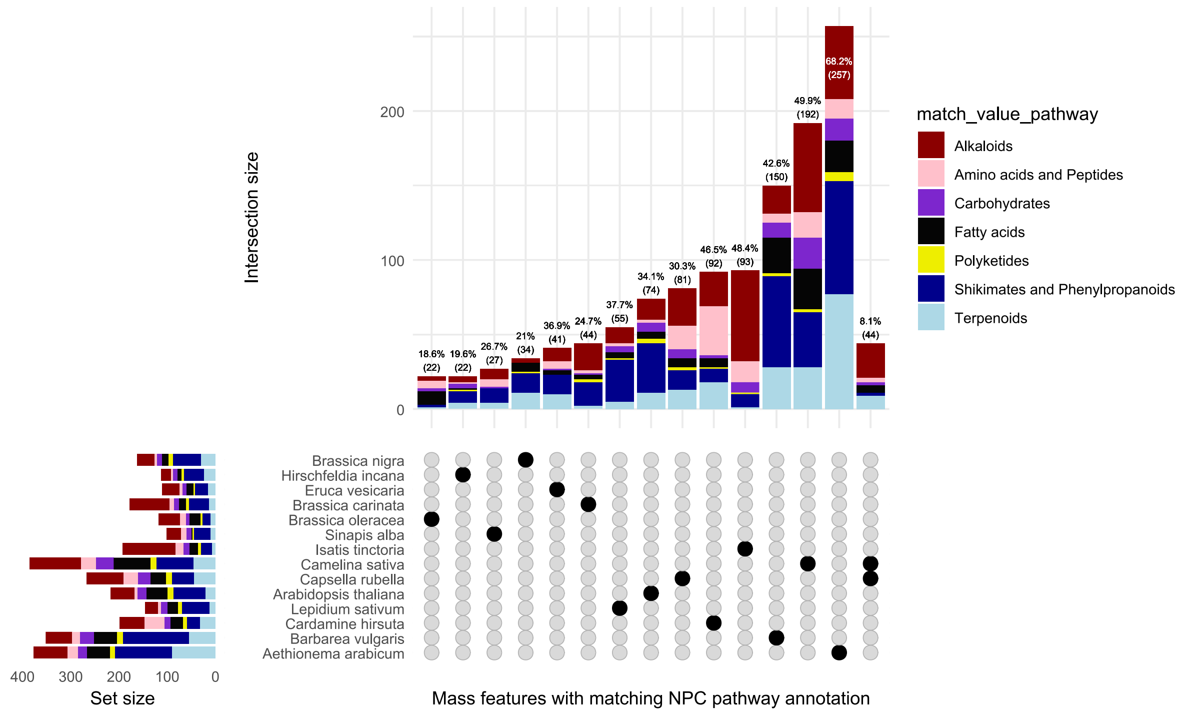


Positive ionization mode, root-specific:

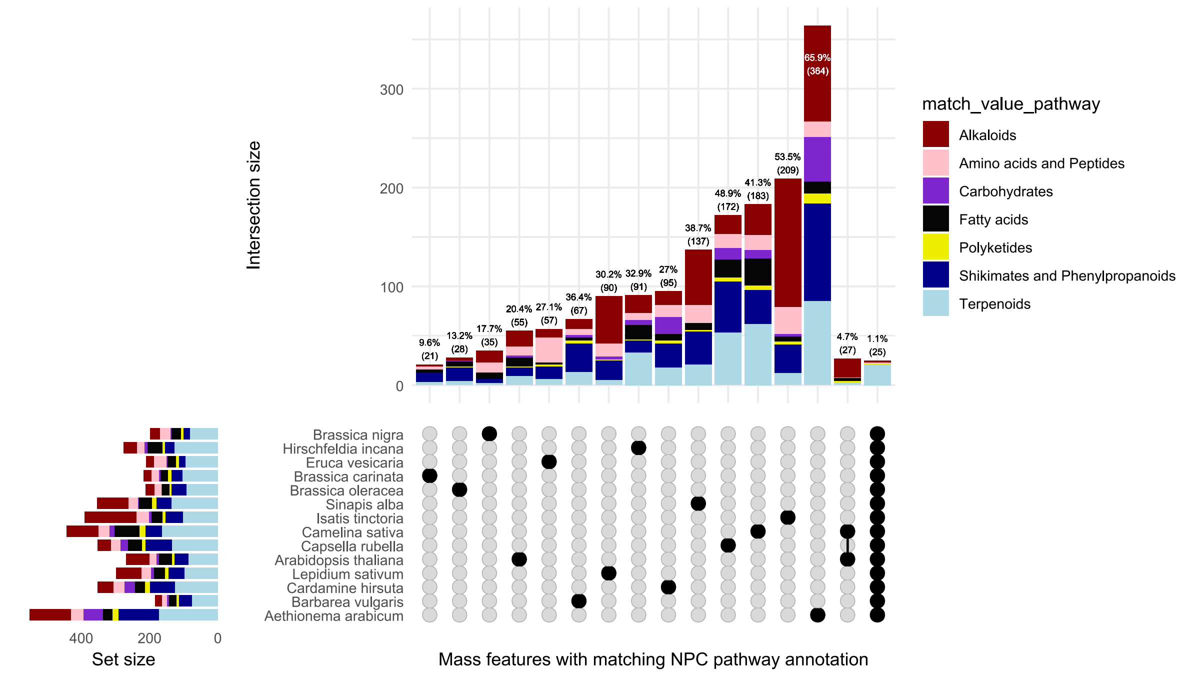

### Negative ionization mode, leaf-specific:

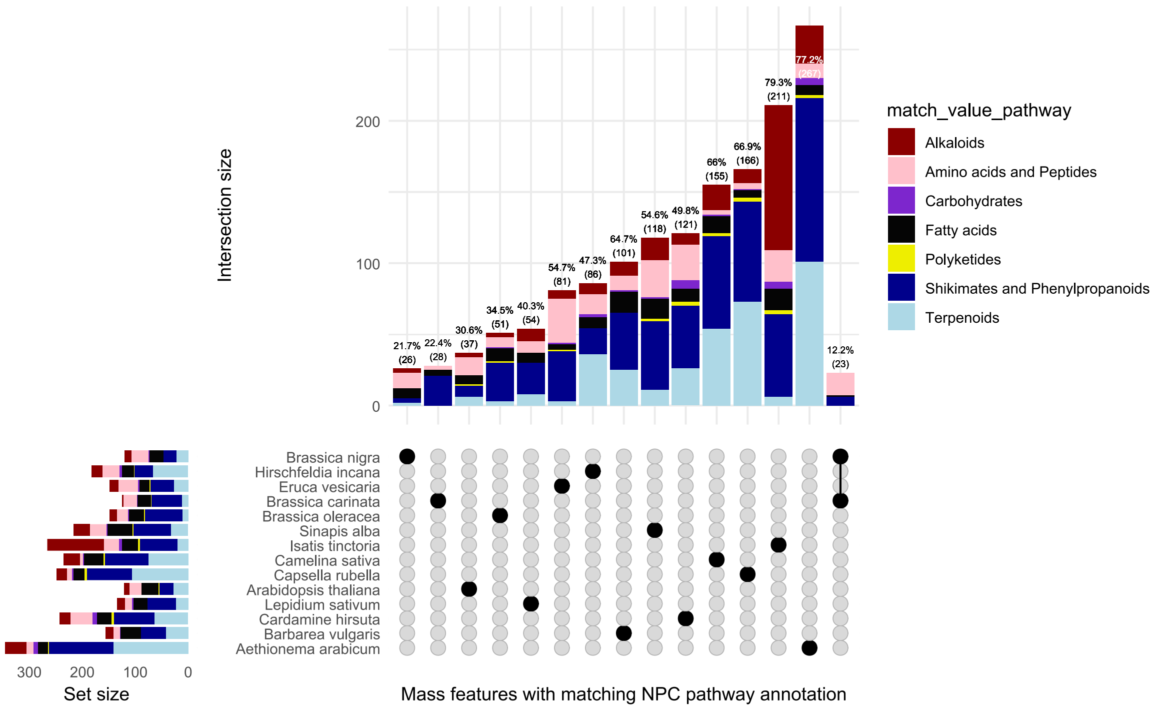


### Negative ionization mode, root-specific:

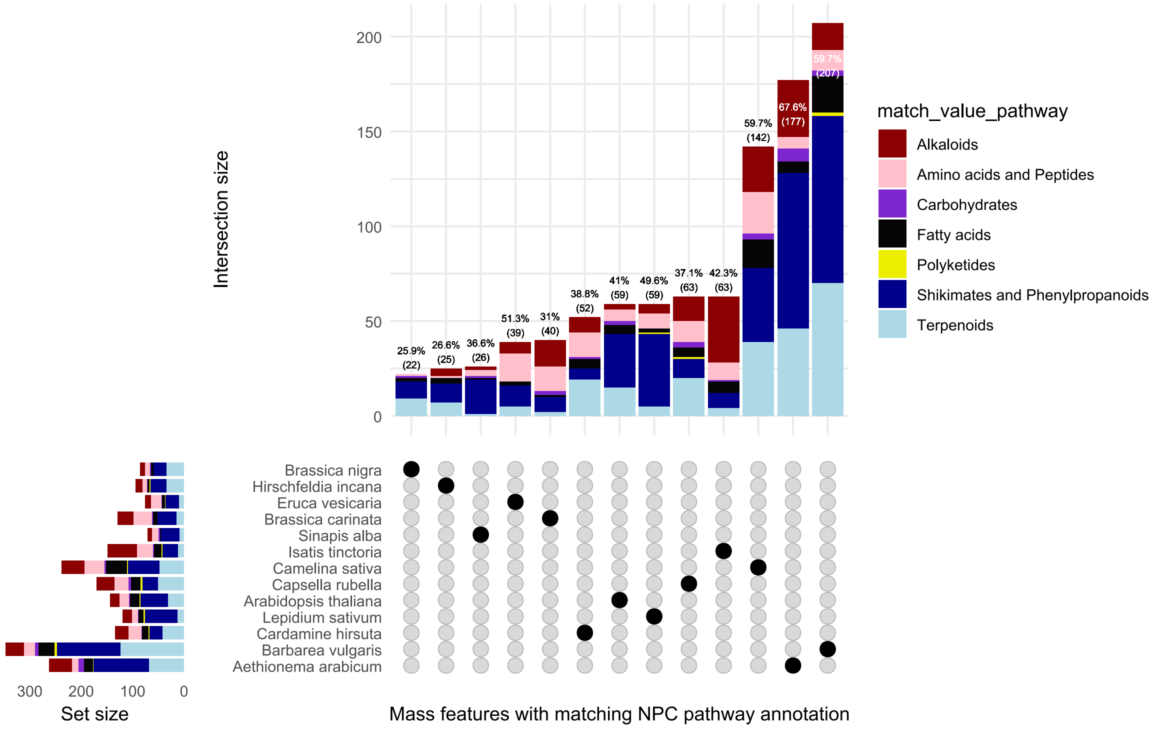


**Fig. S3** Upset plots showing shared and unique annotated metabolite features exclusively found in leaf and root tissue of 14 species.
